# Histone acetylation by SAGA Complex but not by NuA4 Complex is Required for filamentation programme in *Candida albicans*

**DOI:** 10.1101/2025.06.24.661151

**Authors:** Priyanka Nagar, Basharat Bashir Teli, Divya Dinesh, Krishnamurthy Natarajan

## Abstract

*Candida albicans*, a major human fungal pathogen undergoes filamentation from yeast to hyphal state under filamentation-inducing conditions. Gcn5 and Esa1 are key histone H3 and H4 acetyltransferases, respectively, encoded by the budding yeast and other eukaryotes. While Gcn5, a subunit of the SAGA complex, and Esa1, a subunit of the NuA4 complex are critical for *C. albicans* virulence and hyphal induction, how the relative HAT activities impinge on hyphal gene expression during filamentation is less understood. We found that hyphal gene promoters are hyperacetylated at H3K9 and H4 upon filamentation. By creating point mutations in the HAT domain of Gcn5 and Esa1, we investigated the relative requirement of the SAGA and NuA4 HAT activities for filamentation response. We show that Gcn5 HAT activity is essential for hyperacetylation of H3K9 and H4 at promoters and across hyphal gene ORFs. Surprisingly, the Esa1 HAT domain mutation did not impair H4 acetylation at hyphal genes suggesting that Gcn5 HAT activity is sufficient for H4 (and H3K9) acetylation. Paradoxically, the Esa1 HAT mutant formed filaments constitutively and showed elevated H3K9ac and H4ac at promoters under inducing conditions. Furthermore, we show that the basic helix-loop-helix transcriptional regulator Efg1 is essential for Gcn5-mediated hyperacetylation and RNA pol II recruitment to promoters. Thus, our results indicate that the SAGA-mediated H3K9 and H4 acetylation is sufficient and essential for induction of *C. albicans* filamentation.

## Introduction

In eukaryotes, regulation of histone modifications is critical for modulation of gene expression. Acetylation of the histone N-terminal tails leads to an open chromatin structure promoting recruitment and assembly of pre-initiation complex, including RNA polymerase II to gene promoters. SAGA and NuA4 complexes are large, well-characterized histone acetyltransferase complexes (writers), that are composed of the catalytic subunits Gcn5 and Esa1, respectively. Gcn5 is a histone lysine acetyltransferase (HAT) whose activity leads to acetylation of histone H3 at Lys9 and Lys14, and multiple lysine residues in histone H4 (Cai et al. 2011; Cieniewicz et al. 2014).

*Candida albicans* is classified as a critical priority fungal pathogen (WHO fungal priority pathogens list to guide research, development and public health action 2022). The understanding of molecular regulation of virulence and pathogenesis is essential to combat *C. albicans,* which is a widespread opportunistic fungus. A prominent characteristic of *C. albicans* is its ability to switch reversibly between morphogenetic forms-unicellular yeast, pseudohyphal or filamentous hypha form (Whiteway and Bachewich 2007). This unique feature of *C. albicans* species greatly enhances its virulence and survival inside the host (Gow et al. 2011). Various environmental inputs that mimic the microenvironment encountered in human host can lead to morphogenesis of *C. albicans* (Sudbery 2011). Transcriptional regulatory network comprising Efg1, Brg1, Nrg1, Ume6 and Hgc1 control this transition and works in conjunction with the transcriptional regulatory complexes to control hyphal-specific gene expression (Stoldt et al. 1997; Lu et al. 2012; Nobile et al. 2012). In addition to transcription factors, dynamic changes in chromatin structure also modulate gene regulation during hyphal development. Gcn5 and Esa1 are two major histone acetyltransferases (HATs) which have been implicated in histone modification (Sterner and Berger 2000) and have been regarded as essential components of this hyphal transition process in *C. albicans* (Lu et al. 2008; Wang et al. 2013; Chang et al. 2015; Shivarathri et al. 2019). We set out to investigate the role of key histone H3 and H4 acetyltransferases in regulating *C. albicans* pathogenesis, as the regulatory mechanism which interconnects post-translational modifications to pathogenesis of this organism is poorly understood. In this report, we dissected the requirement of the SAGA subunit Gcn5 and the NuA4 subunit Esa1 for H3 and H4 acetylation during the induction of filamentation programme. Our results revealed that both H3K9 acetylation and H4 tetra-acetylation are mediated by Gcn5 and regulate hyphal-specific gene (HSG) expression by stimulation of RNA Pol II (Pol II) occupancy dependent on the Efg1 transcriptional regulator. Our results also revealed, surprisingly, that although Gcn5 is central to the H3 and H4 acetylation at filamentation promoters, the H4 acetyltransferase Esa1 activity keeps the filamentation programme repressed under non-inducing conditions indicating divergent roles for the two major HAT complexes in regulating gene expression and filamentation.

## Materials and Methods

### Strains and growth conditions

*C. albicans* strains were derived from the parental *C. albicans* strain SN95. The various strains were pre-cultured in yeast extract-peptone-dextrose (YPD) for 14–16 h at 30°C, diluted to fresh YPD medium, grown for 4 h with or without 10% FBS (Gibco) at 37°C and harvested. Cell pellets were used immediately or snap-frozen in liquid nitrogen and stored at −80 °C till use.

### Plasmids, strains, and oligonucleotides

All plasmids, strains, and oligonucleotides used are listed in supplementary tables S1, S2 and S3 in the supplemental file.

### Construction of strains and plasmids

Details regarding construction of plasmids and *C. albicans* strains are provided in the supplemental file.

### Whole-cell extract preparation and Western blotting

The cell pellets were resuspended in chilled Winston Buffer (40mM HEPES-NaOH pH 7.5, 350mM NaCl, 10% glycerol, 0.1% Tween 20) with added protease inhibitors (2.5 μg/ml Aprotinin, 2mM Benzamidine, 1mM Dithiothreitol, 2 μg/ml Leupeptin, 2 μg/ml Pepstatin, 100μM PMSF, 10 μg/ml TPCK, 10 μg/ml TLCK), and vortexed in presence of glass beads. The lysates were cleared by centrifugation at 13,000 rpm for 15 min at 4 °C. The proteins were separated on 10% SDS-PAGE gel, transferred onto a Protran nitrocellulose membrane (GE Healthcare) and probed with anti-HA (Roche, 12CA5) or anti-Glucose-6-P-dehydrogenase (Merck, A9521) antibodies detected with ECL-Prime Western blot detection reagent (GE Healthcare) and exposed to X-ray film.

### Histone blots

*C. albicans* cultures were grown in YPD liquid medium till saturation, diluted to fresh YPD and incubated at 30°C in an orbital shaker at 200 rpm. Cells were harvested from 20ml ∼0.8 OD_600_ cultures each by centrifugation and resuspended in 600µl NI buffer (0.25M Sucrose, 60mM KCl, 5mM MgCl_2_, 1mM CaCl_2_ and 0.8% Triton X-100). Cell extracts were prepared by glass bead lysis for 15 min at 4°C, centrifuged at 13000 rpm in a microcentrifuge, and the pellet obtained was resuspended in 500µl 1x SDS-PAGE dye and boiled. Equal volume of samples were loaded in 15% SDS-PAGE gel and blotted to nitrocellulose membrane. The membranes were probed with anti-H3 (Abcam, Ab1791), anti-H3K9ac (Abcam, Ab4441) or anti-H4ac (Millipore, 06-866) antibody.

Preparation of *C. albicans* cells for microscopy.

*C. albicans* cells were harvested and washed with sterile 1× PBS, fixed using 4% (v/v) formaldehyde (prepared in 1× PBS) for 30 min at RT, washed thoroughly with 1× PBS to remove formaldehyde. The cells were then mounted on glass slides and imaged under Nikon Eclipse 90i microscope.

### RNA analysis

Total RNA was isolated from *C. albicans* cells and cDNA synthesized essentially as described before (Singh et al. 2011). Real-time qRT-PCR was carried out in Biorad Real-time PCR system using gene-specific primers and differential expression was calculated by the comparative *C*_T_ method (Livak and Schmittgen 2001). The *SCR1* RNA, an RNA polymerase III transcript was used as endogenous control as described previously (Singh et al. 2011).

### Chromatin immunoprecipitation (ChIP) assay

ChIP assays were conducted essentially as described before (Singh et al. 2011). Briefly, all strains were precultured overnight in YPD medium, grown in fresh YPD medium at 30°C to an OD_600_ of 0.8 or YPD + 10% FBS at 37°C for 2 h for filamentation induction, cross-linked with 1% (v/v) formaldehyde, quenched with 125mM glycine, cells harvested, and chromatin extracts prepared by shearing in Bioruptor (model UCD 300, Diagenode). For chromatin immunoprecipitation, the sheared chromatin extracts equivalent to ∼35 OD_600_ cells was incubated with 2μg of each antibody pre-bound to Dynabeads Protein G (ThermoFisher Scientific, 10004D). The following antibodies were used in ChIP: anti-total H3 (Abcam, Ab1791), anti-H3-K9ac (Abcam, Ab4441), anti-H4ac (Millipore, 06-866), RNA Pol II CTD phospho Ser5 antibody (Abcam, Ab5408) and anti-FLAG (Sigma, F1804).

Immunoprecipitation was carried out for 9 h or 6 h (FLAG IP) at 4 °C, washed, and eluted in 100μl 0.1x TE. The input DNA and immunoprecipitated DNA were purified, diluted (Input DNA 1:10 and IP DNA 1:2), and probed by quantitative real-time PCR for specific regions of interest as well as for the control non-specific regions ca21chr1_1573500–1574000 and *RPS8A* coding regions. For all chromatin immunoprecipitations, enrichment was calculated for target regions and for the control nonspecific region (ca21chr1_1573500–1574000) and specific enrichment calculated with respect to input total chromatin. Additionally, *RPS8A* coding region primers was also included instead of ca21chr1_1573500–1574000 region as an additional non-specific region for an independent confirmation.

### Statistical analysis

The statistical significance of quantitative data was determined by Student’s *t*-test in Graph Pad Prism 6, and a *p*-value ≤ 0.05 was considered statistically significant. The *p*-values are indicated as ∗ (*p* ≤ 0.05), ∗∗ (*p* ≤ 0.01), ∗∗∗ (*p* ≤ 0.001), ∗∗∗∗ (*p* ≤ 0.0001), and ns (*p* > 0.05).

## Results

### Filamentation response elicits histone H3 and H4 acetylation at promoters in vivo

To map the histone acetylation profile during filamentation in *C. albicans*, we induced filamentation by incubating cells in YPD containing fetal bovine serum (FBS; 10% (v/v)) at 37°C for 4h. During this time, the yeast-form cells in YPD produced germ tubes by 1h and formed extensive hyphal cells by 4h (Fig. 1a), as shown previously (Lu et al. 2011). *HWP1* (Hyphal Wall Protein 1) and *ALS3* (Agglutinin-like sequence 3) genes encoding adhesins, required for host cell adhesion, biofilm and pathogenesis (Hoyer et al. 1998; Staab et al. 1999; Zhao et al. 2004; Nobile et al. 2006) are among the highly induced genes during filamentation (Mundodi et al. 2021). Therefore, we reconfirmed the mRNA expression profile of *HWP1* by qRT-PCR analysis and found that *HWP1* was induced rapidly even by 20 min, and by 60 min was induced to ∼150-fold compared to the level in YPD (Fig. 1b). Next, to map the histone acetylation profile during filamentation, we carried out chromatin immunoprecipitation (ChIP) assays using chromatin extracts from WT cells induced in presence of FBS at 37°C for 2h or in extracts from control cells grown at 30°C without FBS, and immunoprecipitated using histone H3K9ac, H4 K5, −8, −12 and −16ac, or H3 antibodies. The DNA was analyzed by qPCR spanning locations upstream of the *HWP1* coding region. The data showed that H3K9ac level, normalized to total H3 level, was induced ∼6-fold and ∼2.5-fold at *HWP1-4* and *HWP1-5* regions, respectively, in FBS compared to YPD alone, but not at *HWP1-3* region (Fig. 1c). We also examined H4 acetylation level by using anti-H4ac antibody, and found that H4ac level was stimulated ∼5-fold and ∼2.5-fold at *HWP1-4* and *HWP1-5* regions, respectively, in FBS compared to YPD alone, but not at the *HWP1-3* region (Fig. 1c). The *HWP1* promoter and the ORF regions showed H3K9 and H4 hyperacetylation under filamenting conditions (Fig. S2a). This significant increase in acetylation showed a similar trend using two different non-specific control regions (Fig. S7a-b). These data indicated that both H3K9 and H4K5, −8, −12 and −16ac levels are stimulated at the *HWP1* promoter region during FBS-induced filamentation.

**Fig. 1.**
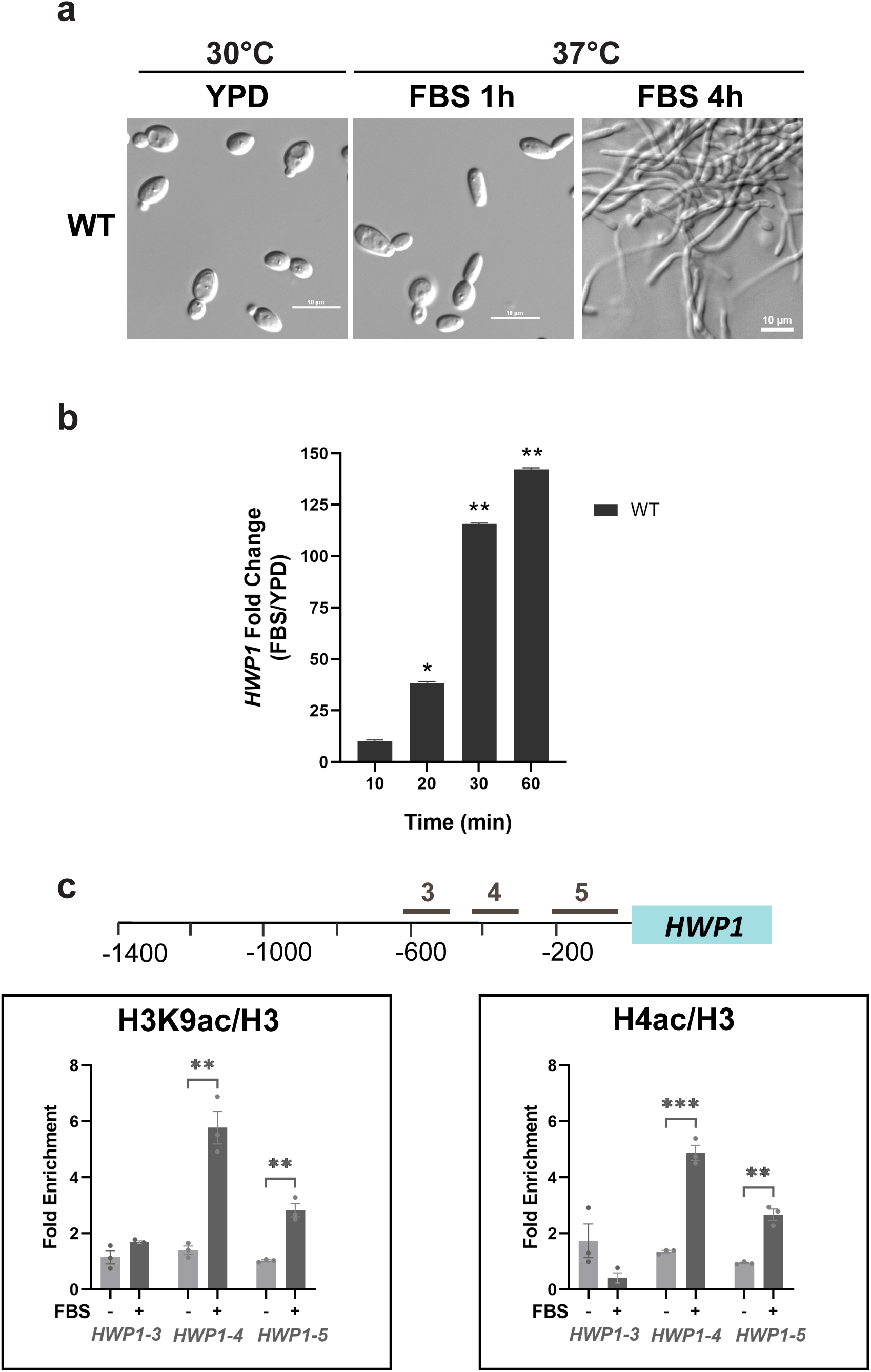
Filamentation-associated histone hyperacetylation elicits hyphal gene expression. (a) Cellular morphology of *C. albicans* under filamentation-inducing conditions. The cells were cultured in YPD alone at 30°C or with 10% (v/v) FBS for 1h or 4h at 37°C, cells harvested, fixed with 4% (v/v) formaldehyde, mounted on glass slides, and images acquired in Nikon Eclipse 90i microscope under 40X objective. The scale bar corresponds to 10μm. (b) mRNA level of hypha-specific gene *HWP1* in wild type cells at different times in YPD + 10% FBS at 37°C. Total RNA was isolated, single-stranded cDNA prepared and qRT-PCR was carried out using SYBR green chemistry. Relative expression level, indicated as fold-change, was calculated as ratio of mRNA level in YPD + 10% (v/v) FBS at 37°C relative to the level in YPD at 30°C. The *C. albicans SCR1* transcript was used as an endogenous control. (c) Chromatin-level histone acetylation at *HWP1* promoter under filamentation condition using ChIP assay. Wild type (WT; SN95) was grown in YPD at 30°C or was induced for filamentation (YPD+10% FBS, 2h, 37°C), and equal amounts of chromatin extracts were used for immunoprecipitation with anti-H3K9ac, anti-H4ac (K5, K8, K12 and K16) antibodies or anti-H3 antibody. The immunoprecipitated DNA was analyzed by qPCR using primer pairs for different regions of *HWP1* promoter, or a Chr I region as a nonspecific control. The data was quantified from three biological replicates each and fold-enrichment plotted. The error bars represent SEM (n = 3). The *p*-values ≤ 0.05 (∗), ≤0.01 (∗∗), ≤0.001 (∗∗∗), and >0.05 (ns) are indicated.

### HAT activity of the SAGA subunit Gcn5 is required for filamentation response

Gcn5 protein contains the histone acetyltransferase (HAT) catalytic domain that carries out histone lysine acetylation by transferring the acetyl moiety of acetyl-CoA to specific lysine residues in histone proteins (Candau et al. 1997). The evolutionarily conserved glutamic acid residue (Glu173) in the *S. cerevisiae* HAT domain was shown to be critical for histone acetylation using *in vitro* histone acetylation assay (Trievel et al. 1999; Kollenstart et al. 2019). Previous studies showed that deletion of *C. albicans GCN5* (*CaGCN5*) or a point mutation Glu188, homologous to Glu173 in *S. cerevisiae* Gcn5, impaired bulk histone H3 acetylation and filamentation (Chang et al. 2015), although the mechanism of how *Ca*Gcn5 HAT activity impacted filamentation at the gene expression level was not clear. Therefore, to first test the expression of Gcn5 protein, we constructed 3xHA-tagged WT and homozygous mutant strains carrying either a single point mutation (Glu188Gln) or as a double mutation (Phe186Ala and Glu188Ala), that are highly conserved across different species (Fig. 2a and Fig. S1a). We examined the expression of Gcn5, Gcn5^FAE-AAA^ and Gcn5^E188Q^ proteins in *C. albicans* whole cell extracts by Western blotting using anti-HA antibody (Fig. 2b). The results showed that while the Gcn5^FAE-AAA^ mutant protein was poorly expressed, the Gcn5^E188Q^ protein was expressed at a substantial level (∼50%) compared to the WT Gcn5 protein in cell extracts from YPD grown cells (Fig. 2c).

**Fig. 2.**
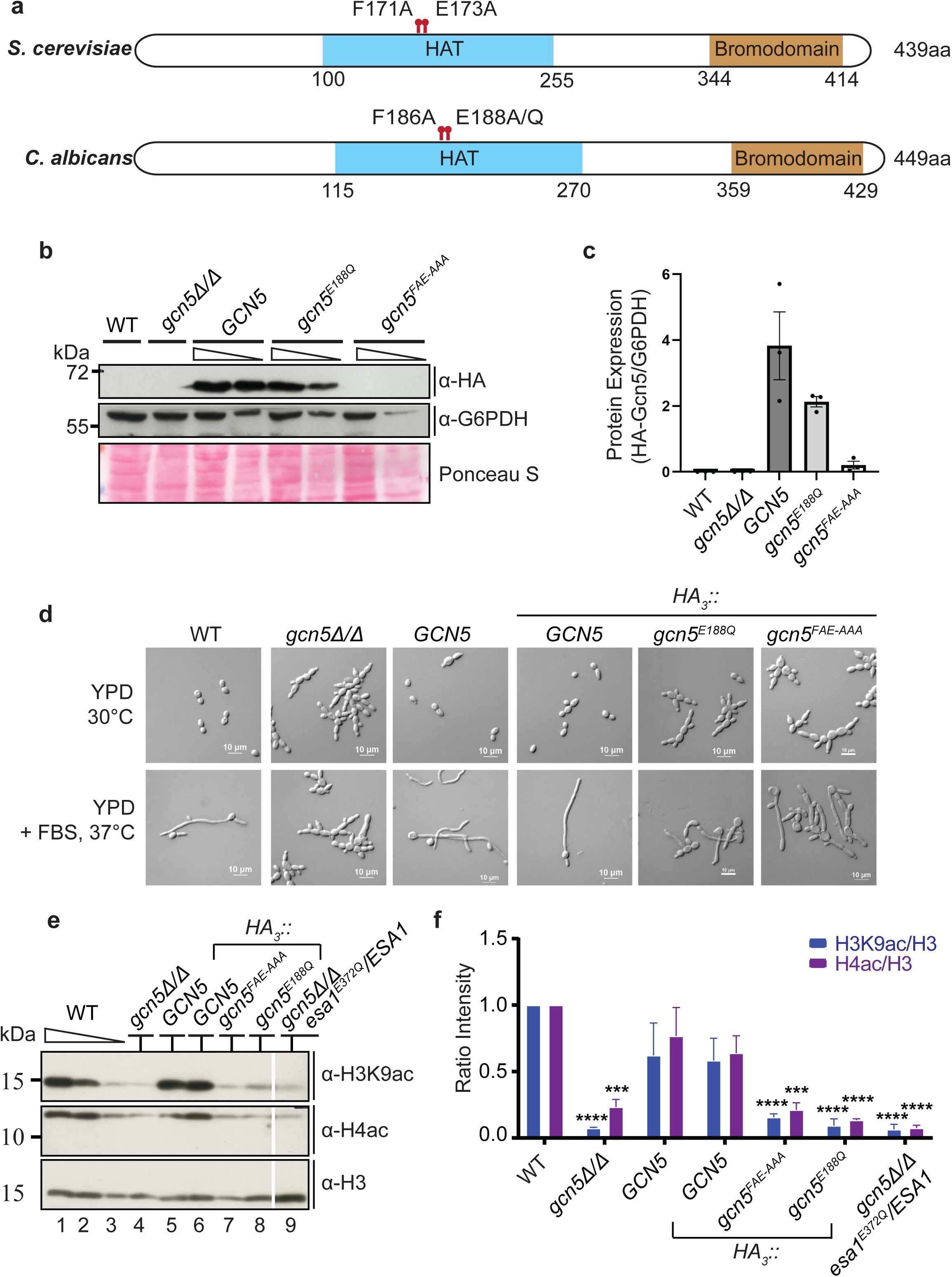
**Gcn5 HAT activity is required for morphogenesis in *C. albicans.*** (a) Schematic representation of *S. cerevisiae* and *C. albicans* Gcn5 amino acid sequences. The HAT domain and the bromodomain regions are shaded, and the catalytic residues in the *S. cerevisiae* and *C. albicans* Gcn5 proteins are also indicated. (b-c) Western blot analysis of WT and mutant Gcn5 proteins. The WT or the catalytically dead Gcn5 single (E188Q), and double (FAE-AAA) mutant strains were cultured in YPD to OD_600_ ∼0.8, cell extracts prepared, and western blot carried out using anti-HA (1:1500), or the anti-G6PDH (1:2500) antibodies, images quantified using Image J and plotted. As controls, untagged WT and the *gcn5Δ/Δ* mutant strains were also used. (d) Cellular morphology of Gcn5 mutants under filamentation-inducing conditions. The strains were cultured in YPD liquid medium with or without 10% FBS and grown for 4h at 30°C or 37°C, cells harvested, fixed with 4% (v/v) formaldehyde, mounted on glass slides, and images acquired in Nikon Eclipse 90i microscope under 40X objective. The scale bar corresponds to 10μm. (e-f) Western Blot analysis of total H3K9ac, H4ac and H3 levels in WT and Gcn5 mutant strains. The mutant and control WT (SN95) strains were grown overnight in YPD liquid media, diluted to fresh YPD and grown for 4h. Lysates were prepared and equal volumes from each sample were loaded on SDS-PAGE, blotted to nitrocellulose membrane and probed with antibodies as indicated. The position of molecular size markers are indicated. The band intensities were quantified using ImageJ and normalized to H3, and plotted as levels relative to that of the WT. The error bars represent SEM (n = 3), and *p*-values ≤ 0.05 (∗), ≤0.01 (∗∗), ≤0.001 (∗∗∗), and >0.05 (ns).

To study the effect of Gcn5 HAT domain mutation on its *in vivo* catalytic activity, we first examined the global H3K9 acetylation level (Fig. 2e). Total histone cell extracts were prepared and resolved on SDS-PAGE gel and probed with different antibodies. The histone blot results revealed that the *gcn5Δ/Δ* mutant had severely reduced bulk H3K9 acetylation level compared to the WT level (Fig. 2e-f). Reintroduction of a copy of either untagged *GCN5* or *HA_3_::GCN5* restored the H3K9ac level, indicating that the addition of 3xHA epitope tag did not alter Gcn5 HAT function. However, both HAT domain point mutants viz., Gcn5^FAE-AAA^ and Gcn5^E188Q^ had severely reduced bulk H3K9 acetylation levels (Fig. 2e-f). We also probed for bulk H4ac level in the same experiment and found that the H4ac level was also substantially reduced in the *gcn5Δ/Δ* and the two Gcn5 HAT domain mutants (Fig. 2e-f). Thus, the evolutionarily conserved HAT domain residues are required for global histone H3K9 and H4 (K5, K8, K12 and K16) acetylation activity in *C. albicans*.

We next examined the filamentation response of the Gcn5 HAT domain mutants Gcn5^FAE-AAA^ and Gcn5^E188Q^ with and without FBS at 37°C for 4h. Whereas the WT cells formed extended hyphae in FBS medium, the *gcn5Δ/Δ* mutant showed constitutive pseudohyphal morphology in both YPD and FBS media (Fig. 2d). The *gcn5^FAE-AAA^* and *gcn5^E188Q^*strains formed constitutive pseudohyphal morphology in YPD, although in FBS medium they formed short hyphae-like structures (Fig. 2d). Overall, the Gcn5 HAT domain mutants showed marked filamentation defect compared to the WT cells, indicating that Gcn5 HAT activity is critical for robust filamentation response in *C. albicans*.

### Gcn5 is required for hyperacetylation of H3 and H4 at hyphal promoters during filamentation

We next tested *HWP1* and *ALS3* mRNA expression after 30 min of filamentation induction in WT, *gcn5Δ/Δ* and *gcn5^E188Q^* mutants. The qRT-PCR data showed that under non-filamentous condition, the *HWP1* and *ALS3* mRNAs levels are not altered by either the *gcn5Δ/Δ* or the *gcn5^E188Q^*mutations with reference to that in the WT strain (Fig. 3a). Under filamentation conditions, the *HWP1* and *ALS3* mRNAs were highly induced (∼20-to 40-fold) by 30 min in the WT strain (Fig. 3b), and their induction was drastically reduced in *gcn5Δ/Δ* or the *gcn5^E188Q^*strains (Fig. 3b). Thus, the loss of *HWP1* and *ALS3* mRNA levels mirror the defective filamentation phenotypes of the *gcn5* mutant strains (Fig. 2d). Therefore, we next decided to study the histone hyperacetylation profiles at these two candidate genes, to investigate whether the decrease in the bulk histone H3K9ac level detected by histone blots (Fig. 2e-f) in the *gcn5* mutants also impacted promoter-level histone acetylation. We define hyperacetylation as enrichment of acetylated histone normalized to total H3 under +FBS and -FBS condition.

**Fig. 3.**
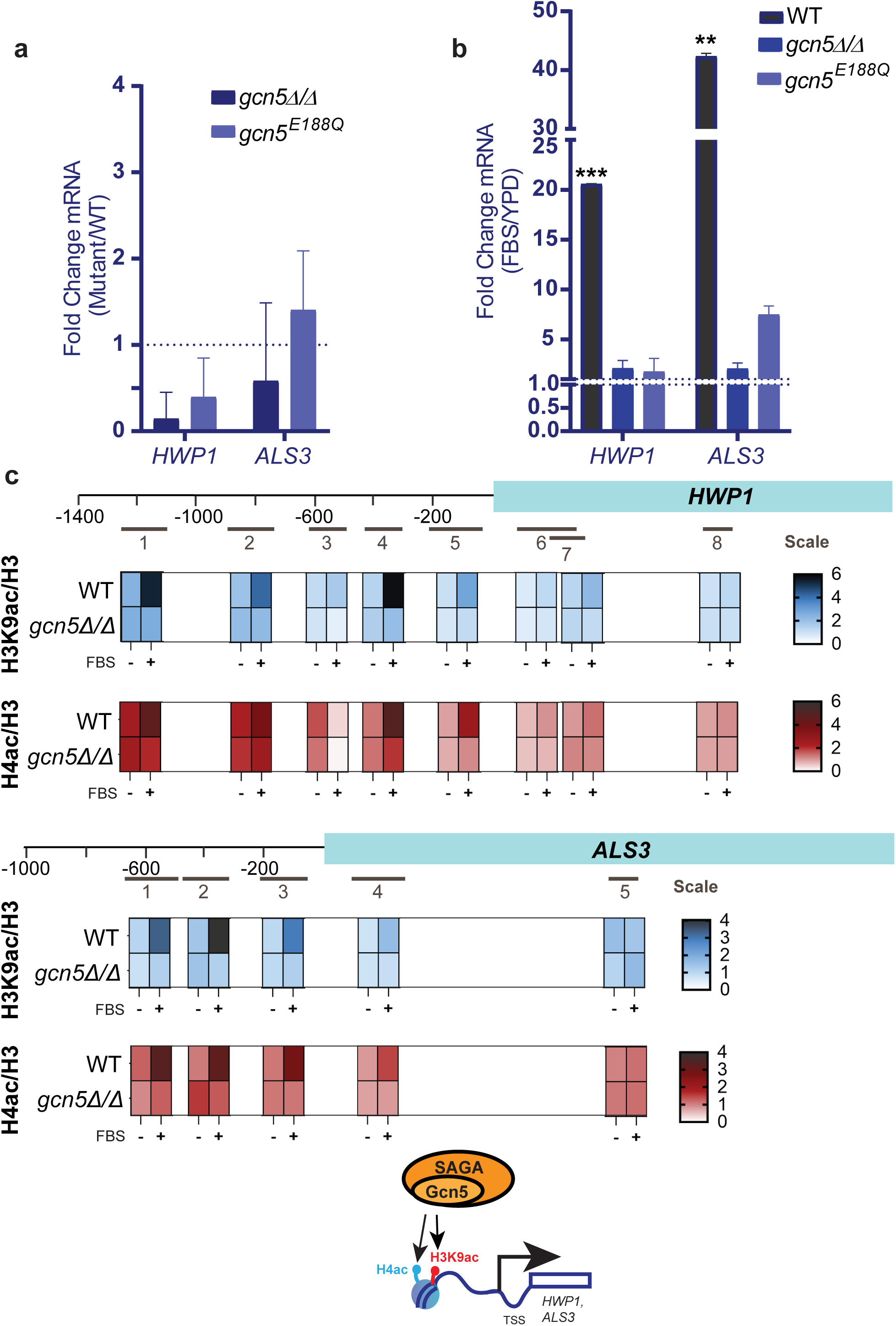
Histone H3K9ac and H4ac at hyphal gene promoters are mediated by Gcn5. (a-b) Hyphal gene expression in wild type and Gcn5 mutants. The mutant and WT strains were grown for 4h in YPD at 30°C, and either induced for filamentation in YPD with 10% FBS for 0.5h at 37°C, or harvested from YPD without induction. Total RNA was isolated, single-stranded cDNA prepared and qRT-PCR was carried out using SYBR green chemistry with primers specific for each of the indicated hyphal genes. Relative expression levels, indicated as fold changes, were calculated either (a) as ratio of mRNA levels in the mutants relative to the levels in WT strain (SN95) in YPD medium at 30°C, or (b) as ratio of mRNA levels in YPD +10% FBS at 37°C relative to the level in YPD at 30°C. The *C. albicans SCR1* transcript was used as an endogenous control. (c) ChIP analysis of chromatin-level histone acetylation at *HWP1* and *ALS3* promoters. Wild type (WT; SN95) and *gcn5Δ/Δ* strains were grown in YPD at 30°C or were induced with 10% FBS for 2h, 37°C, and equal amounts of chromatin extracts were used for immunoprecipitation with anti-H3K9ac, anti-H4ac(K5, K8, K12 and K16) antibodies or the control anti-H3 antibody. The immunoprecipitated DNA was analyzed by qPCR using primer pairs for different regions of *HWP1* and *ALS3* or coding sequences, or a Chr I region as a nonspecific control. The ChIP data was quantified from three biological replicates each, and fold-enrichment was plotted as heatmaps, and the color scale of the enrichment ratio is shown. Note that the ChIP data presented for the WT strain at *HWP1-3*, *-4* and *-5* are the same as in Fig. 1, and experiment conducted along with the other regions *HWP1-1*, *-2*, *-6*, *-7* and *-8* presented here. The thumbnail schematic diagram shows the acetylation of *HWP1* and *ALS3* promoters by Gcn5 subunit of the SAGA complex.

As the bulk H3K9ac levels in the *gcn5Δ/Δ* and the *gcn5* point mutants were comparably reduced, we used the *gcn5Δ/Δ* mutant for the ChIP analysis. Chromatin extracts from WT and *gcn5Δ/Δ* cells incubated with or without FBS were used for immunoprecipitation using histone H3K9ac and H3 antibodies. The ChIP data showed that H3K9ac level was induced ∼2-to 4-fold in FBS in the promoter regions of *HWP1* (*HWP1-1, −2, −4* to *-5*), and ∼3-fold stimulation at *ALS3* (*ALS3-1* to *-3*) in the wild type strain (Fig. 1c, 3c and Fig. S2). At the beginning of the ORF (*HWP1-7* and *ALS3-4*), the H3K9ac level was increased only by about 1.5-fold, whereas in the ORF (*HWP1-8* and *ALS3-5*), the H3K9ac level was not significantly altered under filamentation condition in the wild type strain. The ChIP data further showed that in the *gcn5Δ/Δ* mutant, the H3K9 hyperacetylation was not elevated at the *HWP1* and *ALS3* promoter regions in +FBS compared to -FBS condition (Fig. 3c, Fig. S2).

We next examined histone H4 acetylation in WT and *gcn5Δ/Δ* strains at the same locations in *HWP1* and *ALS3*. The ChIP data showed hyperacetylation of H4 (K5, K8, K12 and K16) at *HWP1* (*HWP1-4* and *-5*) and *ALS3* (*ALS3-1, −2, −3, −4*) promoter regions in the wild-type strain in FBS containing medium. In the *gcn5Δ/Δ* strain, however, the hyperacetylation was impaired (Fig. 3c, Fig. S2), indicating that Gcn5 is the critical HAT mediating H4 (K5, K8, K12 and K16)ac, in addition to its H3K9ac activity at *HWP1* and *ALS3*.

To further assess the correlation between histone acetylation and gene activation, we decided to probe histone acetylation at a control gene not shown to be induced by FBS. Therefore, we examined a previous dataset (Mundodi et al. 2021) for genes with little or no expression change under FBS condition (Serum 37°C/YPD 30°C), and selected *FTR1* gene as the control gene for the ChIP analysis. We carried out qPCR analysis probing for H3K9ac and H4ac with reference to total H3 at the promoter and the coding region of *FTR1.* We found no hyperacetylation of H3K9 and H4ac with reference to H3 under FBS condition in WT strain, normalized to either the Chr I region or the *RPS8A* coding region as controls (Fig. S7c). Thus, the FBS-induced hyperacetylaton at H3K9 and H4 are linked to upregulation of expression of filamentation genes.

A previous report showed that Epl1, a subunit of the NuA4 histone H4 acetyltransferase complex, was recruited to *HWP1* promoter, and H4ac was also shown to be elevated at 30 min of filamentation induction (Lu et al. 2008). These results suggested that the NuA4 recruitment was linked to H4 hyperacetylation, but was not formally tested. Besides, our results showed that H4 acetylation during filamentation was Gcn5-dependent (Fig. 3c).

Therefore, to probe this further, we next decided to test the impact of Gcn5 loss on NuA4 complex recruitment. We constructed FLAG-tagged *EPL1* (*EPL1::FLAG_3_*) in WT and *gcn5Δ/Δ* strains and carried out chromatin immunoprecipitation analysis. Indeed, Epl1 was recruited to the *HWP1* promoter distal region (*HWP1-1*) (Fig. S3a). Interestingly, Epl1 occupancy was comparable at *HWP1-1* under inducing conditions both in the WT and the *gcn5Δ/Δ* strain (Fig. S3a), suggesting that absence of Gcn5 and the attendant loss of histone H3K9 and H4 acetylation did not affect Epl1 occupancy. Taken together, these results showed a central role for Gcn5 in H3K9 and H4 acetylation to induce hyphal gene upregulation.

### HAT activity of CaEsa1 is dispensable for filamentation response

Esa1, the HAT subunit of the NuA4 complex is an essential protein in *S. cerevisiae*, but the Esa1 HAT domain point mutant is viable (Smith et al. 1998; Decker et al. 2008). The *C. albicans ESA1* gene although was shown to be non-essential in a particular study (Wang et al. 2013), a large-scale mutational study reported that *ESA1* is an essential gene (Segal et al. 2018). We attempted to delete *ESA1* in the *C. albicans* SN95 strain but could not obtain a homozygous deletion mutant (data not shown), indicating that *ESA1* is essential in this strain background. Therefore, we decided to create catalytically dead Esa1 mutant by site-directed mutagenesis to probe the role of this HAT in filamentation response. As shown in Fig. 4a, the *C. albicans* Esa1 protein contains an amino-terminal chromodomain and the evolutionarily conserved Glu residue at position 372 in the MYST domain, analogous to the Glu338 in the *S. cerevisiae* Esa1 sequence. Moreover, the *C. albicans* Esa1 protein contains an insert region within the MYST domain that is carboxyl-terminal to the catalytic Glu372 residue.

**Fig. 4.**
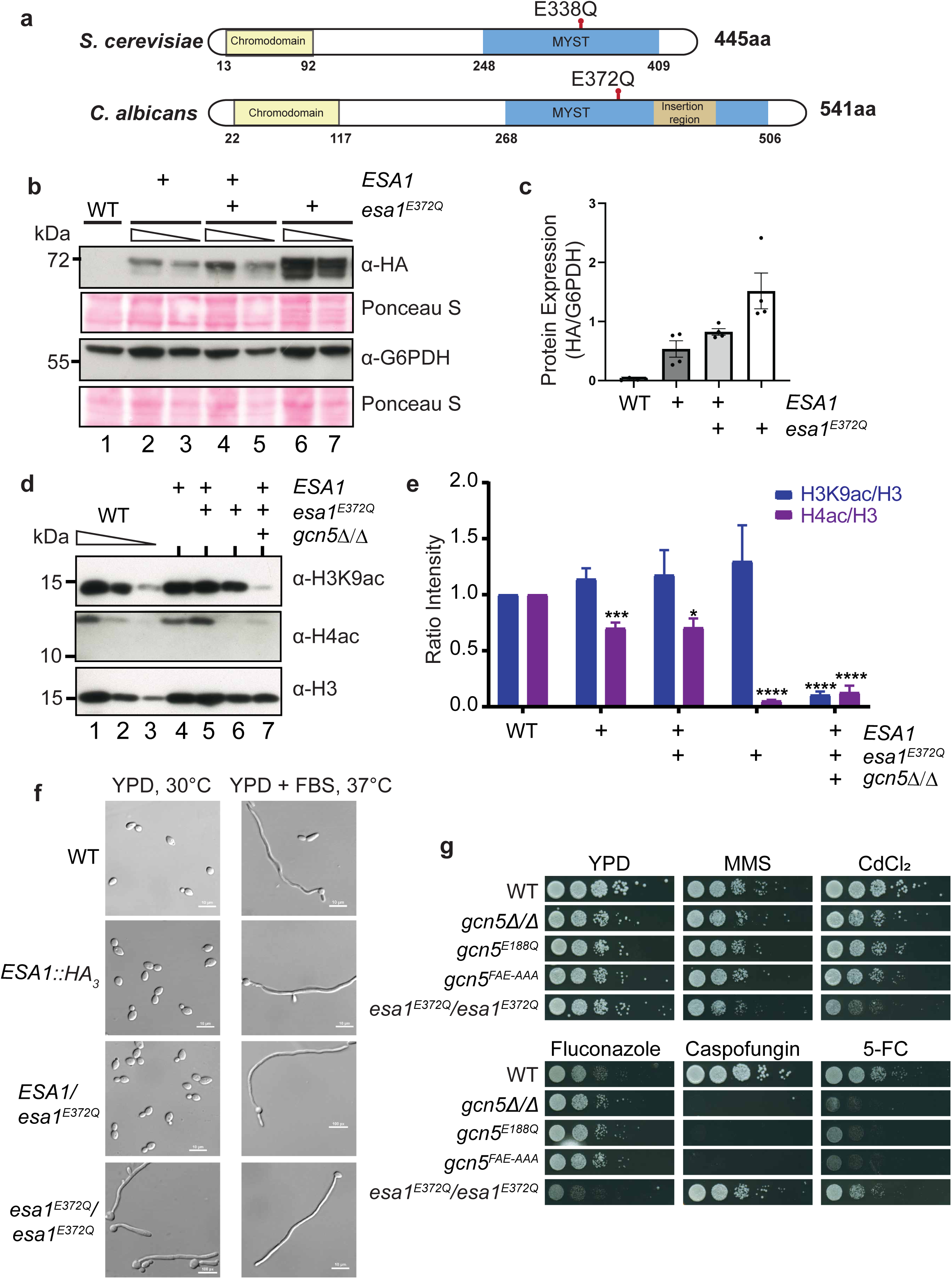
HAT activity of Esa1, a subunit of NuA4 complex is dispensable for filamentation. (a) Schematic representation of Esa1 proteins from *S. cerevisiae* and *C. albicans* showing the evolutionarily conserved catalytic residue in the MYST domain, and the chromodomain. (b) Western blot analysis of Esa1 protein expression. The *ESA1::HA_3_*, heterozygous and homozygous *esa1^E372Q^*mutant strains, and SN95 (untagged control) strain were cultured in YPD at 30°C, cell extracts prepared. Two-fold serial dilutions were analysed by western blot and probed with anti-HA antibody or as control anti-G6PDH antibody. The following strains were used: Lane 1, SN95 (WT); Lanes 2-3, PRI35 (*ESA1::HA_3_/ESA1*), Lanes 4-5, PRI36 (*esa1^E372Q^::HA_3_*/*ESA1*) and Lanes 6-7, PRI40 (*esa1^E372Q^::HA_3_*/*esa1^E372Q^::HA_3_*). (c) Esa1 protein level was quantified from the western blot, normalized to G6PDH level and plotted. (d) Western Blot analysis of H3K9ac, H4ac and H3 levels in the *esa1^E372Q^* mutant strains. The mutant strains and the control SN95 strain were grown in YPD liquid medium for 4h, and cell extracts were analysed by western blotting using anti-H3K9ac, anti-H4ac, and total H3 antibodies. The following strains were used: Lanes 1-3, SN95 (WT); Lane 4, PRI35 (*ESA1::HA_3_/ESA1*); Lane 5, PRI36 (*esa1^E372Q^::HA_3_*/*ESA1*); Lane 6, PRI40 (*esa1^E372Q^::HA_3_*/*esa1^E372Q^::HA_3_*); Lane 7, PRI41 (*gcn5Δ/Δ esa1^E372Q^::HA_3_*/*ESA1*). (e) Quantitation of levels of total H3K9ac and H4ac. The band intensities were quantified using ImageJ, normalized to H3, and plotted as levels relative to that of the WT. The error bars represent SEM (n = 3), *p*-values ≤ 0.05 (∗), ≤0.01 (∗∗), ≤0.001 (∗∗∗), and >0.05 (ns). (f) Cellular morphology of *esa1^E372Q^* mutant. The strains PRI35 (*ESA1::HA_3_/ESA1*), PRI36 (*esa1^E372Q^::HA_3_*/*ESA1*), and PRI40 (*esa1^E372Q^::HA_3_*/*esa1^E372Q^::HA_3_*), and the control WT (SN95) were grown in YPD at 30°C or in YPD+10% FBS at 37°C for 4h. Cells were harvested, fixed with 4% (v/v) formaldehyde, mounted on glass slides and images acquired in Nikon eclipse 90i microscope using 40X objective. The scale bar (10μm) is shown in the bottom right. (g) Growth phenotype of *gcn5* and *esa1* mutants. The WT, *gcn5Δ/Δ*, *gcn5^E188Q^*, *gcn5^FAE-AAA^* and *esa1^E372Q^/esa1^E372Q^*strains were grown in YPD, diluted and serial dilutions spotted on either YPD alone or on YPD with methyl methanesulfonate (0.01%), cadmium chloride (75µM), fluconazole (2.5µg/ml), caspofungin (0.32 μg/ml) and 5-fluorocytosine (30μg/ml). The plates were incubated at 30°C for 36h and images acquired.

We cloned the full-length *C. albicans ESA1* gene bearing 3xHA tag at the 3’ end, and introduced a single point mutation Glu372Gln (Esa1^E372Q^) by PCR-mediated site-directed mutagenesis, and constructed heterozygous and homozygous *ESA1* mutant strains. We first tested the expression level of the WT and the Esa1^E372Q^ mutant proteins by western blot analysis using anti-HA antibody. The results showed that the heterozygous and the homozygous Esa1^E372Q^ mutant proteins are abundantly expressed in *C. albicans* (Fig. 4b-c). To test the effect of Esa1^E372Q^ mutation on bulk histone H4 acetylation, we conducted western blot analysis of H4 acetylation using H4 pan acetyl antibody that detects acetylation at K5, 8, 12 and 16. The western blot results showed that bulk H4ac was severely reduced in the *esa1^E372Q^/esa1^E372Q^* mutant strain, but not H3K9ac that was comparable to the WT level (Fig. 4d-e). The data presented in Fig. 2e, showed that *gcn5Δ/Δ* and the *gcn5* HAT domain mutations substantially reduced H4ac in addition to the H3K9ac level. Therefore, to test if *esa1^E372Q^* mutation further impacted H4ac level in the *gcn5Δ/Δ* background, we attempted to construct homozygous *esa1^E372Q^/esa1^E372Q^*mutation in the *gcn5Δ/Δ* background. However, we were unable to construct this strain even after multiple attempts. Therefore, we used *C. albicans* strain PRI41 bearing heterozygous *esa1^E372Q^/ESA1* mutation in the background of *gcn5Δ/Δ* mutation to examine bulk H3K9ac and H4ac levels. The western blot results showed that in the strain PRI41 (*gcn5Δ/Δ esa1^E372Q^/ESA1*), the bulk H3K9ac level was close to background level (Fig. 4d, lane 7 and Fig. 2e-f), and the bulk H4ac level was further reduced in the indicating that both Gcn5 and Esa1 mediate global H4 acetylation (Fig. 4d-e, Fig. 2e-f).

Next, to examine if Esa1^E372Q^ mutation had any effect on filamentation, the different strains were grown in YPD with or without 10% FBS at 37°C, and cells harvested, fixed with formaldehyde and DIC images were acquired in bright field microscope. Unexpectedly, the *esa1^E372Q^* homozygous strain formed short hyphal structures even at 30°C in YPD (Fig. 4f), and the short hyphae were further elongated in presence of FBS at 37°C. Together, our results showed that Esa1 HAT activity was not required for filamentation in FBS-containing medium, but unexpectedly, loss of Esa1 HAT activity led to constitutive hyphae suggesting that perhaps Esa1 plays an inhibitory role on filamentation in YPD (-FBS) condition. It was previously reported that the HAT domain mutation *esa1^E338Q^* in *S. cerevisiae* led the loss of not only HAT activity, but also genotoxic stress resistance (Decker et al. 2008). Therefore, we tested the *C. albicans* homozygous *esa1^E372Q^* mutant along with the *gcn5* mutants for growth under various stress conditions (Fig. 4g). This comparative analysis showed that the *C. albicans esa1^E372Q^* homozygous mutant, but not the *gcn5* mutants, were impaired for growth in presence of MMS and CdCl_2_ (Fig. 4g). Thus, *C. albicans* showed a requirement for Esa1 HAT activity for genotoxic stress resistance. We also tested growth of *gcn5* and *esa1* mutants in presence of anti-fungal compounds and found that *GCN5* was required for growth in presence of caspofungin and 5-flurocytosine, but not in presence of fluconazole (Fig. 4g). In contrast, *ESA1* was required for growth in presence of fluconazole and 5-flurocytosine, but not in presence of caspofungin (Fig. 4g). Together, these results showed that the Gcn5 and Esa1 HAT activities have overlapping as well as distinct contributions to resistance of *C. albicans* to various stress conditions.

### H4 acetylation at hyphal genes is not dependent on *Ca*Esa1 HAT activity

To determine the requirement of Esa1 HAT activity for histone acetylation and hyphal gene expression, we first measured the mRNA levels of *HWP1* and *ALS3* with and without induction by FBS. The qRT-PCR data showed that the homozygous *esa1^E372Q^*mutation led to ∼35-to 70-fold upregulation of *HWP1* and *ALS3* even in YPD without FBS with respect to the WT cells (Fig. 5a). In FBS medium, while the expression of *HWP1* and *ALS3* was upregulated ∼20-to 40-fold in the WT strain, no upregulation was found in the *esa1^E372Q^*mutant (Fig. 5b). Thus, the strong upregulation of *HWP1* and *ALS3* mRNAs in the *esa1^E372Q^* mutant in YPD and YPD plus FBS, is correlated to the filamentous growth phenotype (Fig. 4f).

**Fig. 5.**
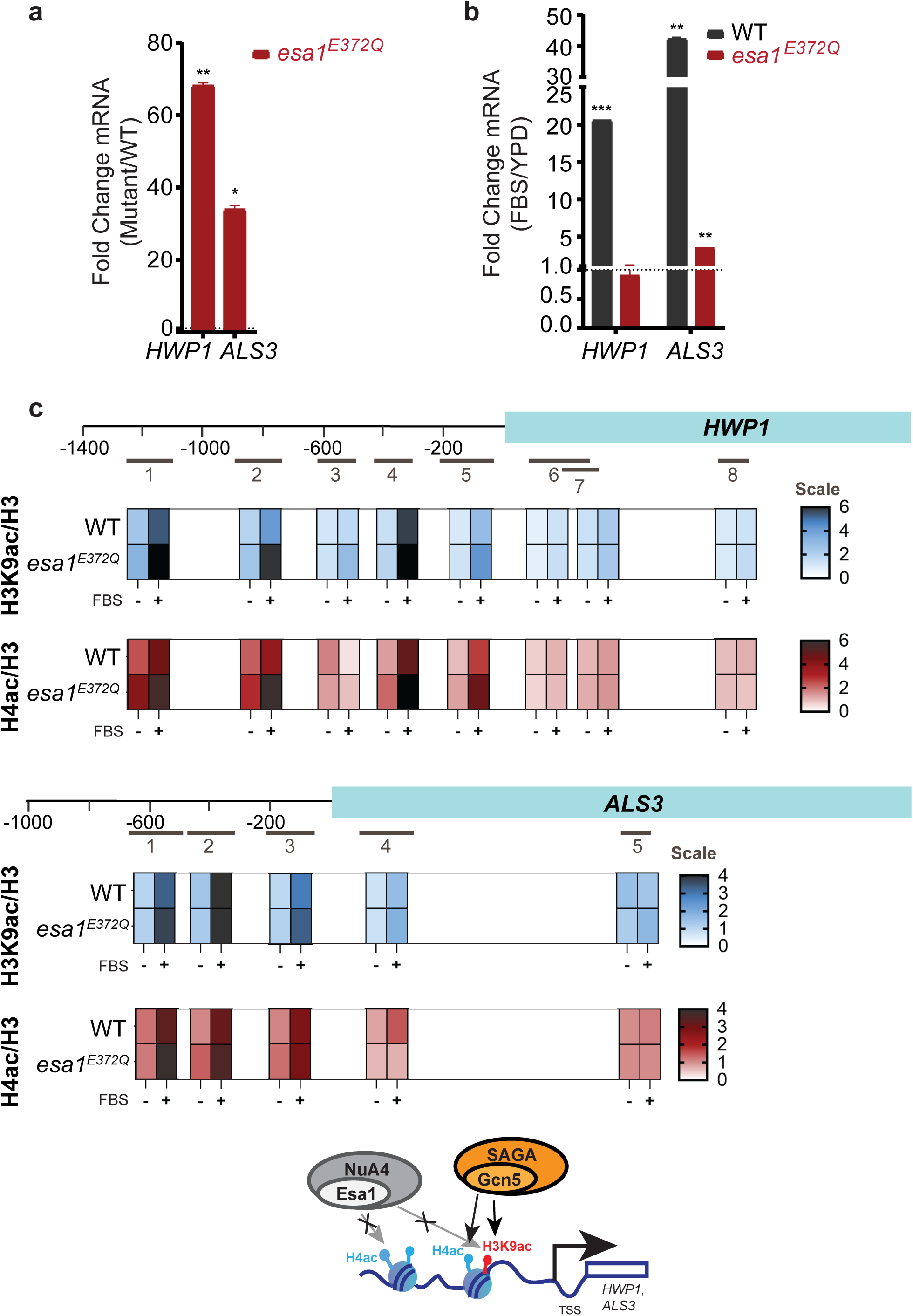
H4ac at hyphal genes is not dependent on *Ca*Esa1 HAT activity. (a-b) Hyphal gene expression in wild type and *esa1^E372Q^* (PRI40) mutant. The mutant and WT strains were grown for 4h in YPD at 30°C, and either induced for filamentation in YPD with 10% FBS for 0.5h at 37°C, or harvested from YPD without induction. Total RNA was isolated, cDNA prepared and qRT-PCR was carried out using SYBR green with primers specific for *HWP1* and *ALS3* transcripts. Relative expression levels, indicated as fold-change, were calculated as the ratio of (a) the mRNA levels in the mutant strains relative to the levels in the WT strain (SN95) in YPD medium at 30°C, or (b) mRNA level in YPD with10% FBS at 37°C relative to the level in YPD at 30°C. *C. albicans SCR1* transcript was used as an endogenous control. (c) ChIP assay to measure the level of histone H3K9 acetylation and histone H4 acetylation at *HWP1* and *ALS3* promoters in wild-type (WT; SN95) and *esa1^E372Q^* (PRI40) strains. Cells were grown in YPD at 30°C or induced for filamentation (YPD+10% FBS, 2h at 37°C), cross-linked with formaldehyde, harvested, lysed, and chromatin fragmented by sonication. Equal amounts of chromatin extracts were used for immunoprecipitation with anti-H3K9ac, anti-H4ac or, as control, anti-H3 antibodies and the immunoprecipitated DNA was analyzed by qPCR using the indicated primer pairs for *HWP1* and *ALS3* genes or a Chr I region as a nonspecific control. The location of the primers is shown in the schematic diagrams. The data was quantified from three biological replicates each and fold-enrichment was plotted as heatmaps at *HWP1* and *ALS3* in WT and *esa1^E372Q^* strains, and the color scale of the enrichment ratio is shown on the right.

We next investigated the levels of histone H3 and H4 acetylation in the *esa1^E372Q^* mutant by ChIP assays. Chromatin extracts from formaldehyde cross-linked cells were prepared and immunoprecipitated using anti-H4ac, -H3K9ac and -H3 antibodies. The enriched DNA was quantified by qPCR using primers for the indicated locations in the *HWP1* and *ALS3* genes (Fig. 5c). The ChIP data showed that H4ac level increased by at least 2-to 3-fold at the promoter locations (*HWP1-4* and *-5*) in the wild type strain upon induction of filamentation (Fig. 5c, Fig. S4). At *ALS3* (*ALS3-1, −2, −3, −4*), H4 hyperacetylation was induced in the WT strain upon filamentation. In the *HWP1-6, −7, −8* and *ALS3-5* ORF regions, we found no stimulation of the H4 acetylation by FBS in the WT strain.

Next, we analyzed H4ac levels at different locations in the *HWP1* and *ALS3* genes in the *esa1^E372Q^* mutant. Surprisingly, we found that the induction of H4ac was not impaired in the *esa1^E372Q^* mutant at the *HWP1* and *ALS3* promoter regions (Fig. 5c, Fig. S4). Together, these data showed that the H4 acetylation at *HWP1* and *ALS3* promoters was not dependent on Esa1 during induced expression under filamentation conditions, although the bulk, cellular-level H4 acetylation was highly dependent on Esa1 as shown in the histone blots (Fig. 4d-e). Next, we analyzed H3K9ac in WT and in the *esa1^E372Q^*chromatin extracts. The ChIP data revealed hyperacetylation of H3K9ac at the *HWP1* (*HWP1-1, −2, −4* to *-5*) and at *ALS3* (*ALS3-1, −2, −3, −4*) promoter in the WT strain, and was essentially unaffected by the *esa1^E372Q^* mutation (Fig. 5c; Fig. S4). Moreover, the H3K9ac at *HWP1* (*HWP1-6, −7, −8*) and *ALS3* (*ALS3-4, −5*) ORFs in the WT strain was also comparable to that in the *esa1^E372Q^* mutant. Thus, H3K9ac and H4 (K5, K8, K12 and K16)ac at *HWP1* and *ALS3* during filamentation is not dependent on Esa1, indicating the primacy of Gcn5 for H3 and H4 acetylation during filamentation (Fig. 3 and 5).

### Efg1 is required for SAGA-mediated histone acetylation during filamentation

Next, we wanted to examine the role of Gcn5 and Esa1 in regulating the expression levels of transcription factors controlling filamentation. The regulatory network controlling filamentation involves the induction of the transcriptional activators *BRG1* and *UME6* expression, and the repression of negative regulator *NRG1* to initiate and sustain filamentation (Kadosh and Johnson 2005; Chow et al. 2021). In addition to these transcriptional regulators, environmental signals through different signal transduction pathways like MAPK and cAMP/PKA pathways converge on the key transcription regulatory factor Efg1 (Stoldt et al. 1997). Although Efg1 is essential for initiation of filamentation, Ume6 is required for hyphal extension (Banerjee et al. 2008; Carlisle et al. 2009). The negative regulator Nrg1 downregulates hypha-specific gene expression thereby acts as a checkpoint for progression of filamentation. Thus, we first analyzed the expression levels of *BRG1, UME6, NRG1* and *EFG1* in cells grown in YPD at 30°C and found that the expression was not altered in either *gcn5* deletion or the *gcn5* point mutant (Fig. 6a). Interestingly, in the *esa1^E372Q^*mutant, *BRG1* and *UME6* were upregulated in YPD medium, while the expression of *EFG1* seem to be partially down-regulated (Fig. 6a). Consistent with the elevated *BRG1* and *UME6* mRNA levels, the *esa1^E372Q^* mutant cells produced hyphae constitutively (Fig. 4f). Under filamentation conditions, the transcriptional upregulation of *BRG1* and *UME6* was impaired in the *gcn5* mutants and the *esa1^E372Q^* mutant. Together our mRNA expression data suggested that Gcn5 and Esa1 HAT activities are required for expression of the filamentation transcriptional regulators as well.

**Fig. 6.**
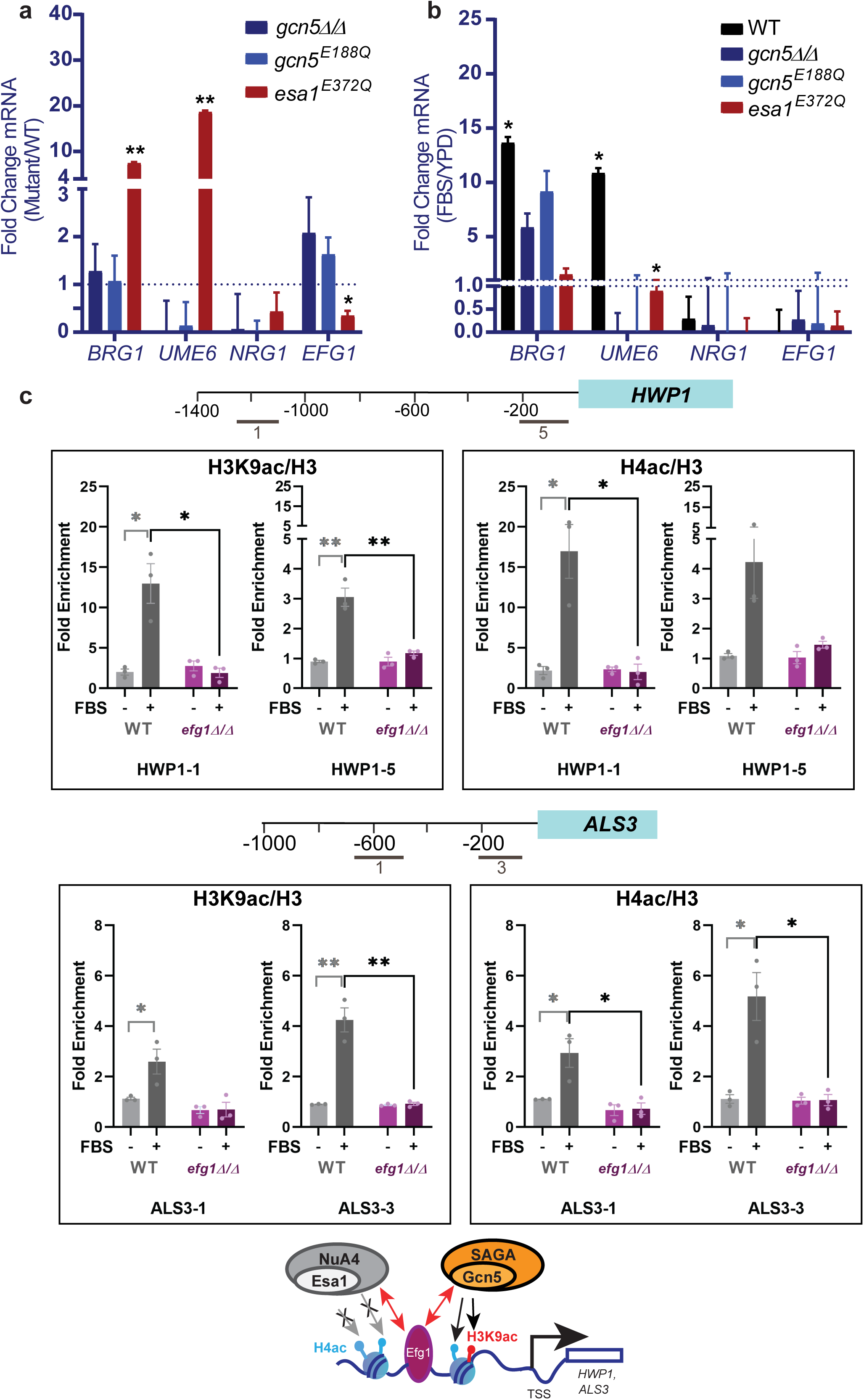
Efg1 is required for SAGA-mediated histone acetylation. (a-b) Hyphal gene expression in wild type, *gcn5Δ/Δ*, *gcn5^E188Q^* and *esa1^E372Q^* (PRI40) mutants. The mutant and WT strains were grown for 4h in YPD at 30°C, and either induced for filamentation in YPD with 10% FBS for 0.5h at 37°C, or harvested from YPD without induction. Total RNA was isolated, cDNA prepared and qRT-PCR was carried out using SYBR green with primers specific for *BRG1, UME6, NRG1* and *EFG1* transcripts. Relative expression levels, indicated as fold-change, were calculated as the ratio of (a) the mRNA levels in the mutant strains relative to the levels in the WT strain (SN95) in YPD medium at 30°C, or (b) mRNA level in YPD with 10% FBS at 37°C relative to the level in YPD at 30°C. *C. albicans SCR1* transcript was used as an endogenous control. (c) ChIP assay was used to measure histone H3K9 acetylation and histone H4 acetylation at *HWP1* and *ALS3* promoters in wild type (WT; SN95) and *efg1Δ/Δ* strains. Cells were grown in YPD at 30°C or induced for filamentation (YPD+10% FBS, 2h at 37°C) conditions, and chromatin immunoprecipitation carried out with anti-H3K9ac, anti-H4ac or as control anti-H3 antibodies. The immunoprecipitated DNA was analyzed by qPCR using primer pairs for *HWP1* and *ALS3* or a Chr I region as a nonspecific control. The data was quantified from three biological replicates each and the error bars represent SEM (n = 3). The *p*-values ≤ 0.05 (∗), ≤0.01 (∗∗), ≤0.001 (∗∗∗), and >0.05 (ns) are indicated.

Next, we asked if Efg1 has a role in Gcn5/SAGA complex-mediated histone acetylation. We carried out H3K9ac and H4ac ChIP analysis at *HWP1* and *ALS3* promoters in the wild type and *efg1Δ/Δ* strains. The ChIP data showed that the hyperacetylation of H3K9 and H4 (K5, K8, K12 and K16)ac in WT cells at *HWP1* and *ALS3* promoters was severely reduced in the *efg1Δ/Δ* mutant (Fig. 6b-c). Thus, Efg1 is essential for Gcn5/SAGA-mediated H3K9 and H4 acetylation of hyphal gene promoters during filamentation response.

### RNA Polymerase II recruitment at HSGs is dependent on Gcn5-mediated acetylation

Next, we asked if RNA polymerase II recruitment is dependent on the acetylation function of Gcn5 or Esa1 at hyphal gene promoters. We carried out ChIP assays using anti-Pol II antibody using chromatin extracts from WT, *gcn5Δ/Δ* and *esa1^E372Q^*strains; and Pol II occupancy was assessed at *HWP1* and *ALS3* genes (Fig. 7a) by qRT-PCR. In WT strain, we found strong stimulation of Pol II occupancy by ∼2-to 4-fold in the core promoter regions (*HWP1-5* and *ALS3-3*) in FBS compared to YPD alone (Fig. 7b). In the *HWP1-6, −7, −8* and *ALS3-4, −5* representing the ORF regions, the Pol II occupancy was further enhanced (∼10-to 15-fold) in the WT strain (Fig. 7b). In the *gcn5Δ/Δ* mutant, however, the Pol II occupancy was completely impaired, down to the level in YPD condition at all regions examined (Fig. 7b). Interestingly, in the *esa1^E372Q^* strain, the Pol II occupancy was largely unaffected in ORF (*HWP1-6, −7, −8*) (Fig. 7b), or only partially reduced (*ALS3-4, −5)*, indicating that Esa1 activity is largely dispensable for Pol II recruitment at these genes. We next asked if the Pol II recruitment is Efg1-dependent. Therefore, we carried out ChIP analysis in WT and *efg1Δ/Δ* strains. The ChIP data showed that Pol II occupancy at the *HWP1* and *ALS3* genes was completely abrogated in the *efg1Δ/Δ* strain (Fig. 7c), indicating a tight requirement of Efg1 for Pol II recruitment to the promoters examined. Overall, we conclude that Pol II recruitment to the filamentation promoters and ORF region is dependent on Efg1 and Gcn5, but not Esa1.

**Fig. 7.**
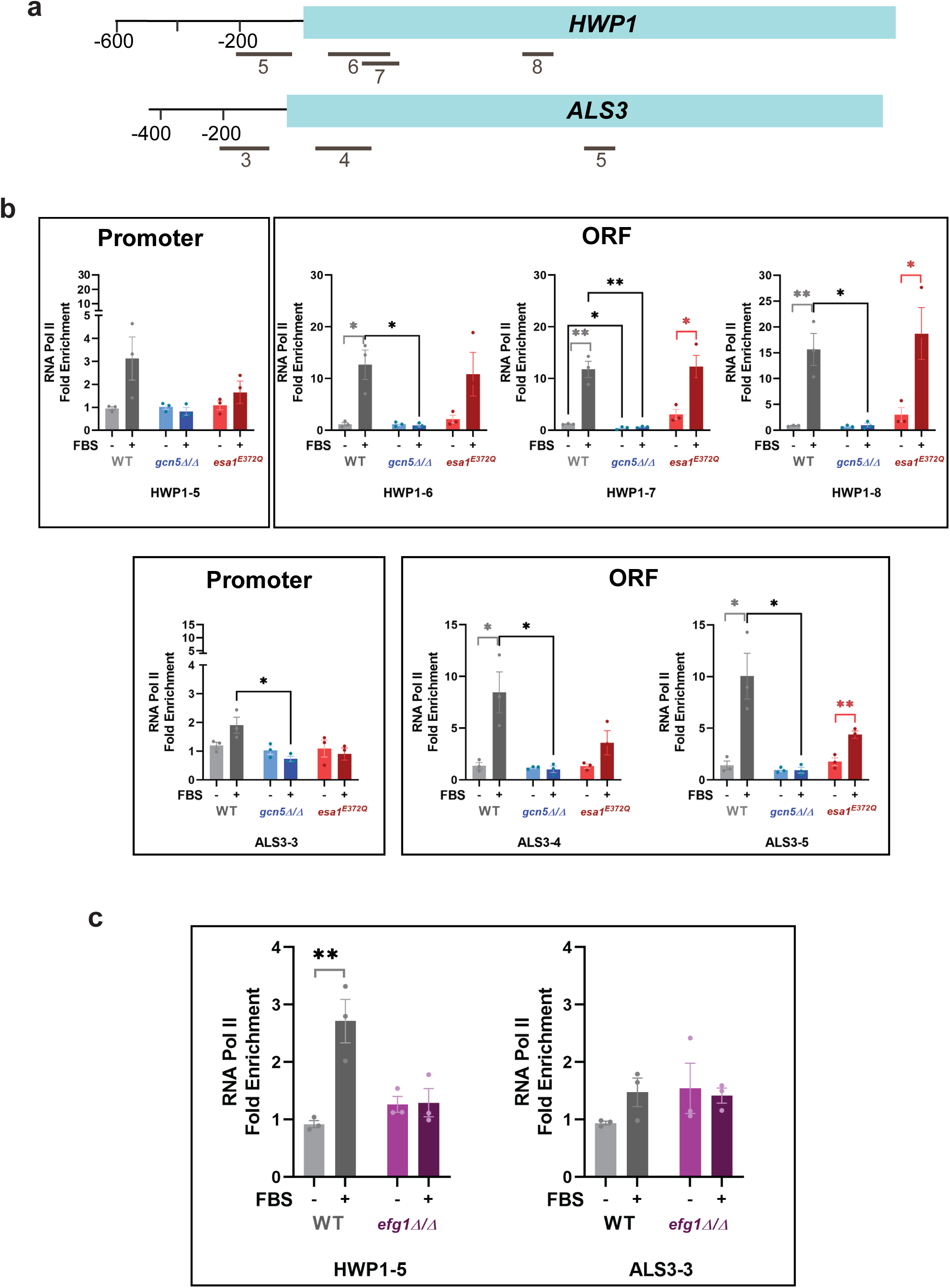
RNA Polymerase II recruitment at hyphal genes is dependent on Gcn5-mediated acetylation. (a) Schematic map of *HWP1* and *ALS3* genes. (b-c) Pol II recruitment was analyzed by ChIP assay at *HWP1* and *ALS3* promoters in (b) the wild-type (WT; SN95), *gcn5Δ/Δ*, *esa1^E372Q^/esa1^E372Q^*and (c) *efg1Δ/Δ* mutant strains. Cells were grown in YPD at 30°C or induced for filamentation (YPD+10% FBS, 2h at 37°C) conditions, and chromatin immunoprecipitation was carried out with equal amounts of chromatin extracts using antibody against Pol II CTD (Ser5P). The Pol II enrichment was normalized to Chr I region (non-specific control) and the data from three biological replicates was plotted as fold-enrichment of specific region with reference to the background. The error bars represent SEM (n = 3) and the *p*-values ≤ 0.05 (∗), ≤0.01 (∗∗), and >0.05 (ns) are indicated.

## Discussion

We have identified a central role for Gcn5 histone acetyltransferase, a subunit of the SAGA complex, for filamentation in *C. albicans*. We found that Gcn5 HAT activity is required for hyperacetylation of histone H3K9 and H4 (K5, K8, K12 and K16), and high level transcriptional induction of filamentation genes *HWP1* and *ALS3* during filamentation response. The SAGA complex subunits Gcn5 and Ubp8 activate hyphal formation in *C. albicans* (Zhu et al. 2021). In contrast, loss of Spt8 results in derepression of hyphal gene expression and filamentous phenotype (Rashid et al. 2022; Teli et al. 2024). These results suggest both activating and repressive functions for SAGA complex in *C. albicans* filamentation. Thus, interaction and functional association of SAGA complex modules determine the cellular morphology of *C. albicans*.

*ESA1* encodes histone H4 acetyltransferase and is an essential gene in *S. cerevisiae* (Smith et al. 1998; Clarke et al. 1999), as well in other fungi (Gómez et al. 2008; Soukup et al. 2012), drosophila (Zhu et al. 2007) and mouse (Hu et al. 2009). However, a previous study reported that the *C. albicans ESA1* is not essential for vegetative growth in strain BWP17, although the *esa1Δ/Δ* mutant showed decreased global histone H4 acetylation level, and could not form hyphae due to impaired hyphal gene expression (Wang et al. 2013). However, we could not obtain *esa1Δ/Δ* mutant in *C. albicans* strain SN95 using multiple deletion strategies (data not shown) suggesting that the different *C. albicans* strain backgrounds employed in the two studies could have led to contrasting phenotypic outcomes. Therefore, we constructed *C. albicans* strain bearing homozygous Esa1^E372Q^ (Glu372Gln) mutation that led to loss of bulk H4 acetylation but showed WT growth (Fig. S1d). Interestingly, the *esa1^E372Q^* mutation showed constitutive filamentation even without inducing conditions (Fig. 4f), and led to elevated levels of hyphal gene expression (Fig. 5a-b).

Given that our data showed a strong Gcn5-dependent H4 acetylation at the filamentation gene promoters, we examined if the Esa1 HAT activity impacted promoter-level H4 acetylation during filamentation. Surprisingly, the Glu372Gln (Esa1^E372Q^) mutation in the Esa1 catalytic domain resulted in loss of bulk histone H4 acetylation but did not impact Esa1 protein level and promoter-level H4 acetylation (Fig. 4 and 5). The Esa1^E372Q^ mutation also led to deregulated expression of *BRG1* and *UME6* and in turn *HWP1* and *ALS3* hyphal genes under non-inducing conditions. Thus, the Esa1 H4 acetylation activity is dispensable for induction of filamentation response and the attendant gene expression, indicating that the Esa1 HAT activity could be negatively controlling filamentation in the absence of filamentation signals. The transcriptional circuitry controlling filamentation is intricately laced with several transcriptional regulators as summarized in a schematic model (Fig. 8). Briefly, in yeast cells, Nrg1 down-regulates the expression of the transcriptional activator *UME6* and hyphal-specific *HWP1* and *ALS3* genes by binding to their promoters. Thus Nrg1 activity maintains cells in yeast form by suppressing filamentation (Lu et al. 2011). However, upon induction of filamentation, Nrg1 expression is downregulated leading to Brg1 binding to promoters and activation of target genes including that of *UME6* (Banerjee et al. 2008; Cleary et al. 2012; Lu et al. 2012; Martin et al. 2013). Efg1 is a critical transcription activator of filamentation (Lu et al. 2008), and under hypoxic conditions functions as a repressor (Desai et al. 2015). In yeast-form cells, Efg1 remains bound to *HWP1* and *ALS3* gene promoters (Lu et al. 2008; Lassak et al. 2011). While *EFG1* mRNA expression is largely comparable between non-filamentous (YPD) and filamentous conditions (serum, 37°C), *EFG1* mRNA translation is repressed in serum at 37°C (Desai et al. 2018; Mundodi et al. 2021). Consequently, Efg1 level is diminished in hyphal cells. Paradoxically, *EFG1* deletion led to non-filamentous cells indicating that Efg1 has an essential role in filamentation.

**Fig. 8.**
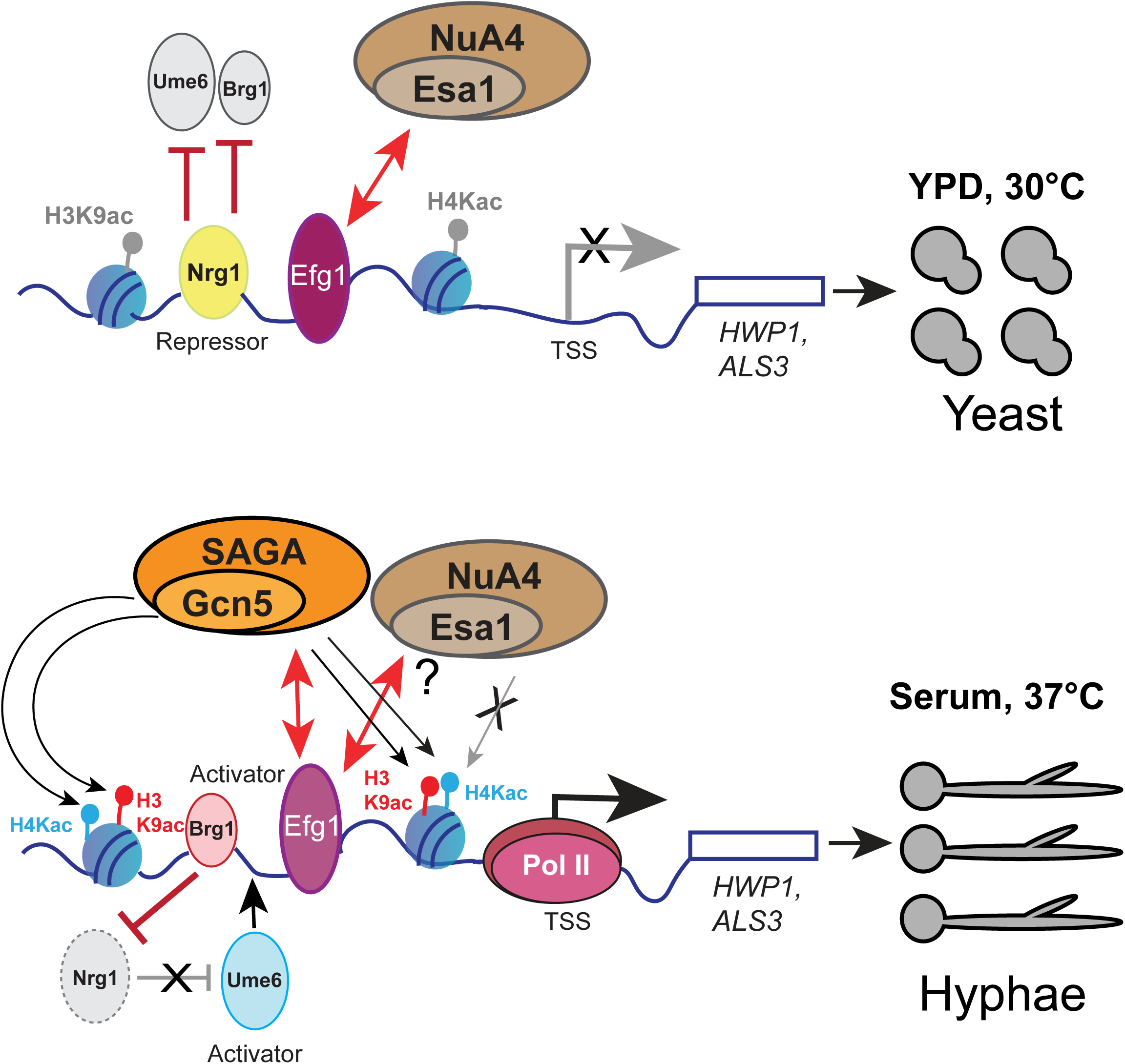
Model showing the role of SAGA and NuA4 HAT complexes in hyphal-specific gene activation. Wild-type *C. albicans* grow as yeast-phase cells in YPD at 30°C, where the hyphal genes such as *HWP1* and *ALS3* are largely repressed; in FBS medium at 37°C, however, the expression of *HWP1* and *ALS3* are strongly induced and the cells form hyphae. Upon filamentation induction, Gcn5, the HAT subunit of SAGA, carries out hyperacetylation of histone H3K9 and histone H4 (K5, K8, K12 and K16) at the promoters of *HWP1* and *ALS3* genes. In *gcn5Δ/Δ* mutant, both H3K9 and H4 (K5, K8, K12 and K16) acetylation is significantly reduced at the promoters, and the cells form pseudohyphae constitutively in YPD, and failed to form hyphae upon induction in FBS. Although Esa1, the HAT subunit of NuA4 complex, is the predominant H4 acetyltransferase for global H4 acetylation, at *HWP1* and *ALS3* promoters, the H4 acetylation was unaffected by the catalytically dead *esa1^E372Q^* mutation. Remarkably, the *esa1^E372Q^* mutation led to strong upregulation of *HWP1* and *ALS3* in YPD leading to constitutive filamentation suggesting that Esa1/NuA4 complex is required to maintain yeast-form cells in YPD by limiting histone acetylation at the promoters. The SAGA-mediated histone hyperacetylation is dependent on the transcriptional regulator Efg1 supporting the model that Efg1 recruits the SAGA complex to modify chromatin in a manner leading to enhanced pol II occupancy during filamentation.

Our results have shown that under filamentation condition, Efg1 is required for SAGA/Gcn5-mediated acetylation of the *HWP1* and *ALS3* genes. Interestingly, Gcn5 also acetylated H4 (K5, K8, K12 and K16) over these chromatin regions. Moreover, deletion of *EFG1* also led to impaired Pol II occupancy at the core promoter *in vivo*. While NuA4 is recruited to filamentation promoters in an Efg1-dependent manner (Lu et al. 2008), our results surprisingly showed no requirement of Esa1 for H4 acetylation to induce filamentation. On the contrary, the Esa1^E372Q^ HAT mutant formed filaments constitutively (Fig. 4f), indicating that NuA4 recruitment to filamentation gene promoters aids in maintenance of chromatin state in a manner that permits filamentation only under appropriate filamentation-inducing conditions. It is unclear how NuA4/Esa1 functions to maintain chromatin state at these gene promoters. As our ChIP results showed no dependence on Esa1 for H4ac at the HSGs examined (Fig. 5), it is conceivable that Esa1 activity could be required for targeted acetylation at certain other promoters such as *UME6* and *BRG1* that are upregulated in the Esa1^E372Q^ mutant. Esa1 could have non-histone related acetylation activity as reported for autoacetylation of Esa1, and acetylation of Eaf1 and Yng2 subunits of NuA4 complex in *C. albicans* (Lu et al. 2011; Wang et al. 2018). To our knowledge, this is the first report demonstrating a critical requirement for the Gcn5 HAT activity for H3 and H4 acetylation to induce filamentation. Thus, the two HAT complexes participate in the same pathway, where Esa1/NuA4 inhibits hyphal gene expression under YPD condition, while Gcn5/SAGA induces gene expression when filamentation cues are provided. Thus, *C. albicans* employs the two predominant HAT enzyme complexes, SAGA and NuA4 in distinct ways for homeostatic morphological states dependent on the induction signals. Our results suggest that modulating HAT activity, for example by small molecules, could be an attractive strategy for controlling of *C. albicans* filamentation and virulence.

## Data Availability

Supplemental material containing figures and methods are available at GENETICS online; all strains and plasmids are available upon request.

## Author Contributions

PN and KN conceptualized the study and analysed the data. PN, BBT, and DD performed the experiments. PN and KN analysed data and drafted the manuscript. K.N. acquired funding, supervised the project and finalized the manuscript.

## Author Notes

Conflict of Interest: The author(s) declare no conflict of interest.

## Supporting information

Legends for Supplental Figures

## Supplementary Figures and Tables

**Fig. S1.** Gcn5 and Esa1 are involved in regulating growth and filamentation of *C. albicans*.

**Fig. S2.** Histone H3K9ac and H4ac at hyphal gene promoters is mediated by Gcn5.

**Fig. S3.** NuA4 complex recruitment is independent of Gcn5 at HSGs.

**Fig. S4.** *Ca*Esa1 HAT activity is not required for H4 and H3K9 acetylation at HSGs.

**Fig. S5.** Construction of *HA_3_::gcn5^E188Q^* complemented strain expressed from native promoter.

**Fig. S6.** Construction of *ESA1::HA_3_* and *esa1^E372Q^*::*HA_3_* plasmid.

**Fig. S7.** Gcn5-mediated acetylation is linked to gene expression and filamentation.

**Table S1.** List of plasmids

**Table S2.** List of strains

**Table S3.** List of oligonucleotides

