## Supplemental Fig S1-S7 for "Histone acetylation by SAGA Complex but not by NuA4 Complex is Required for filamentation programme in *Candida albicans*"

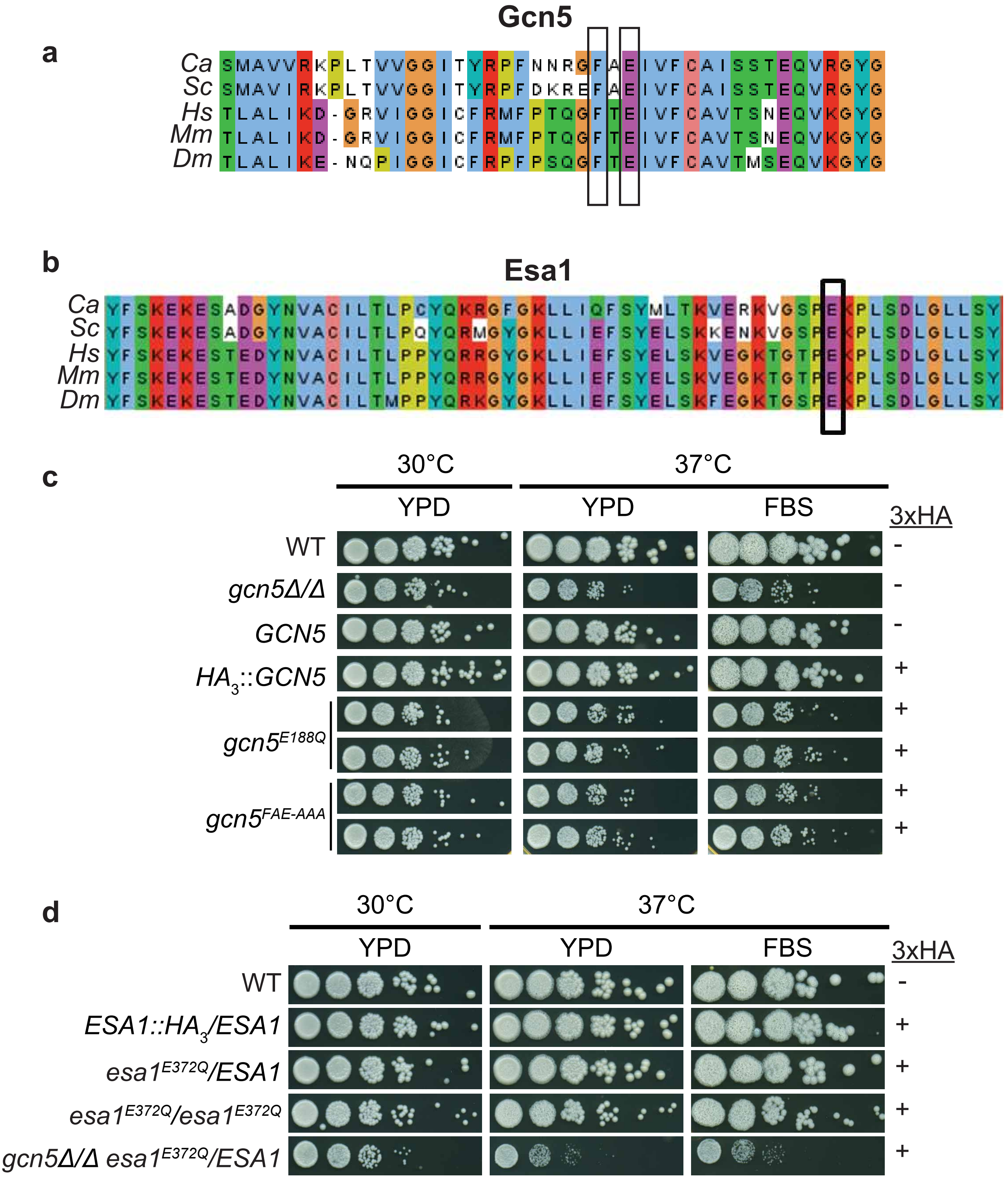


Nagar et al., Fig. S1

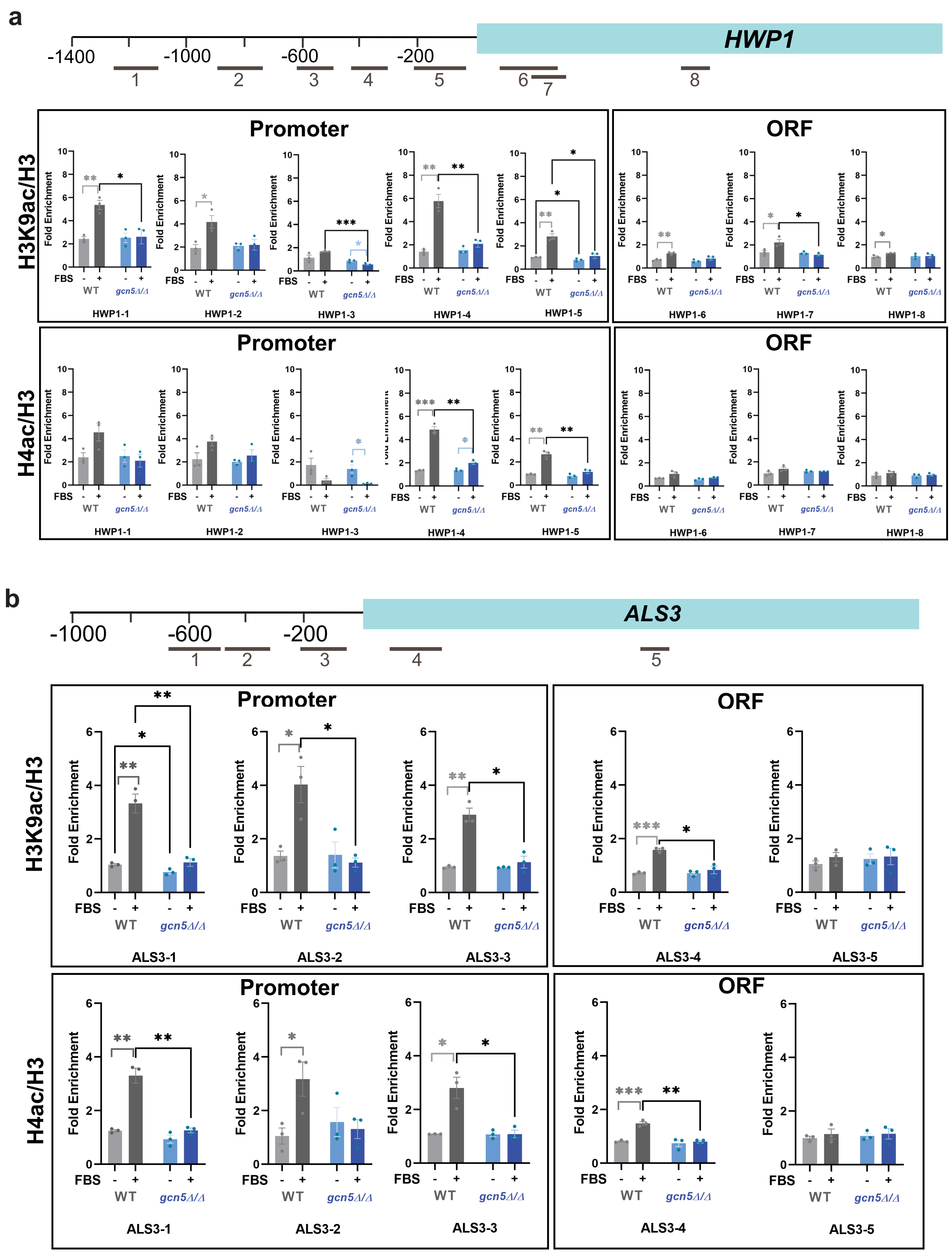


Nagar et al., Fig. S2

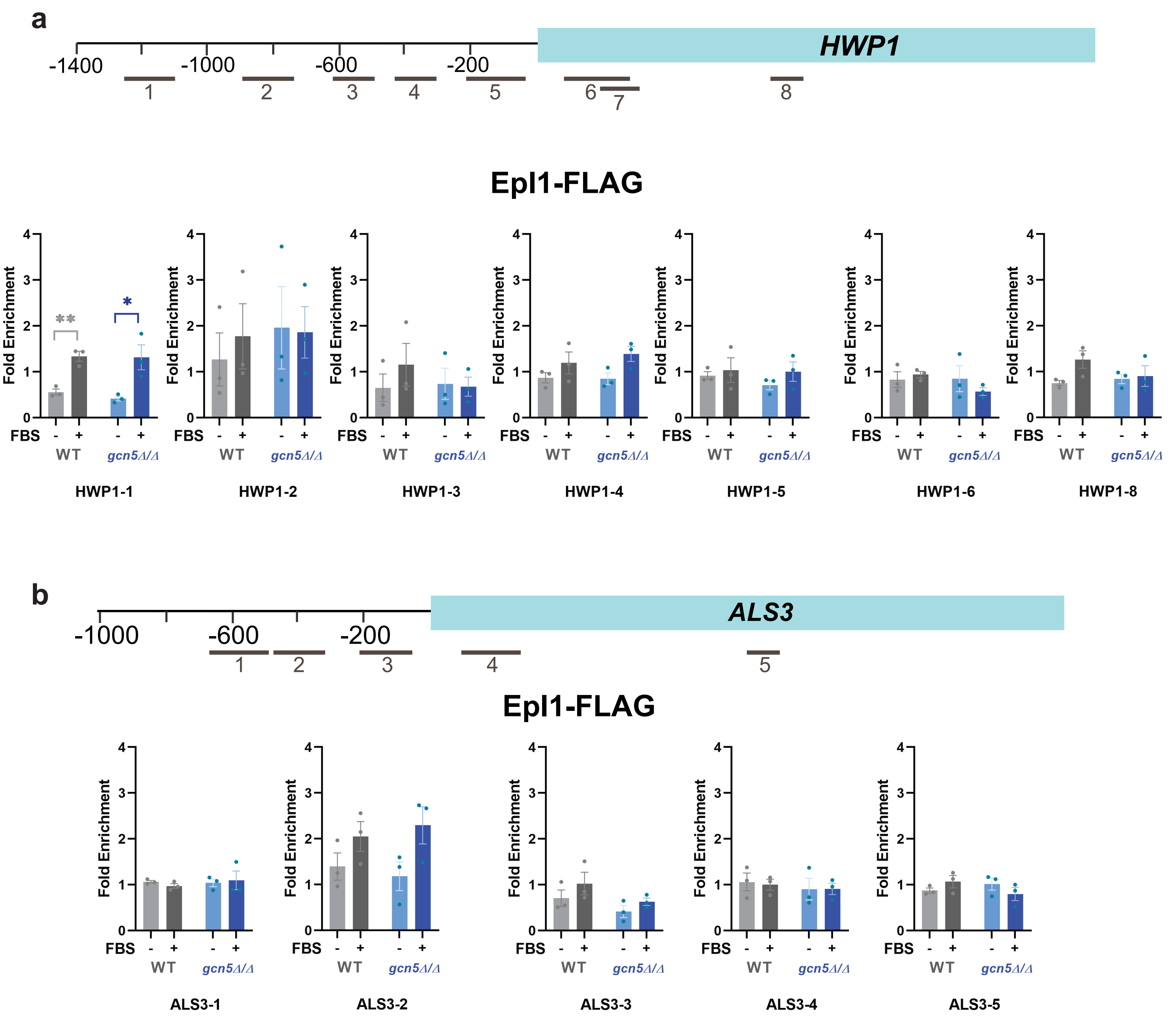


Nagar et al., Fig. S3

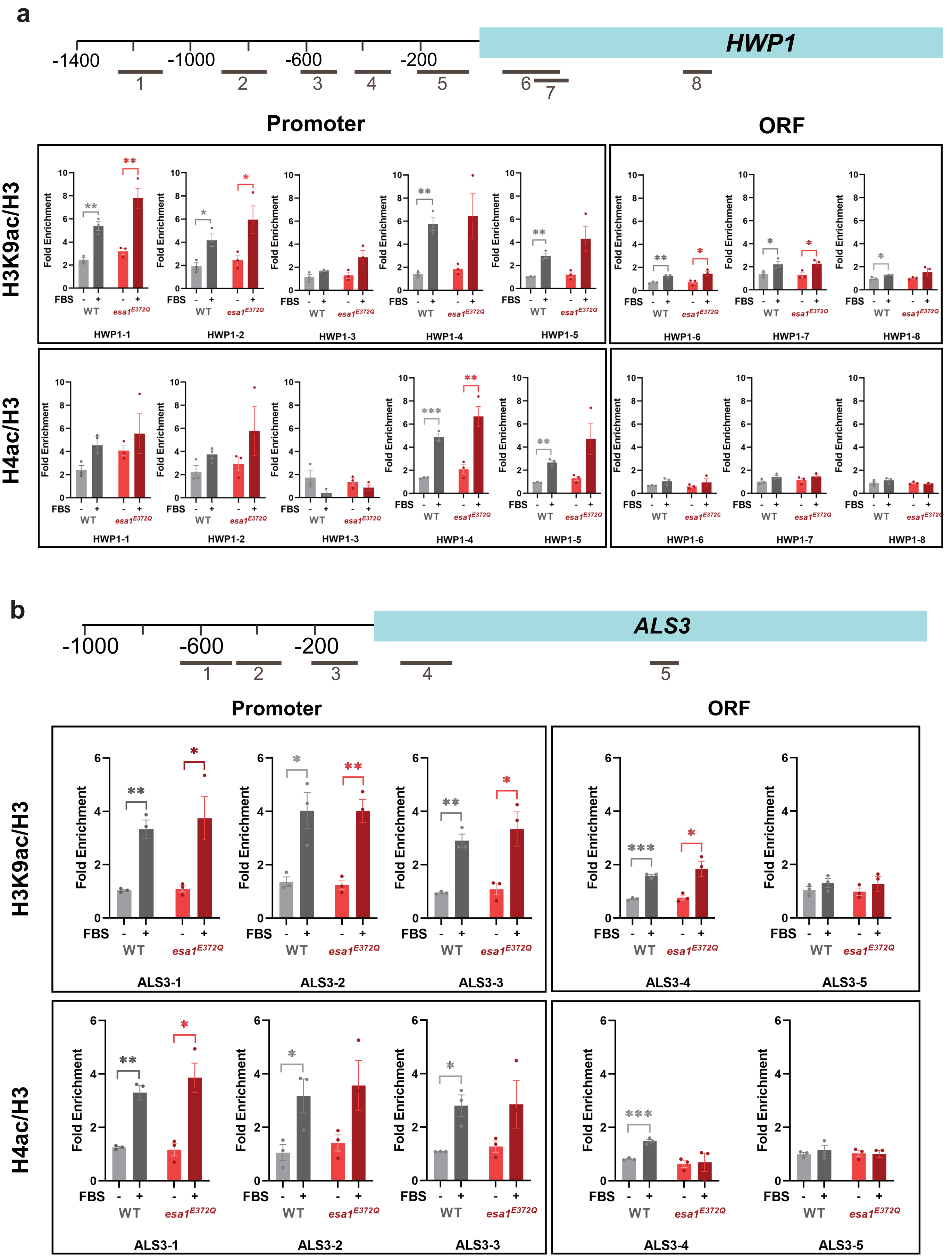


Nagar et al., Fig. S4

**a**

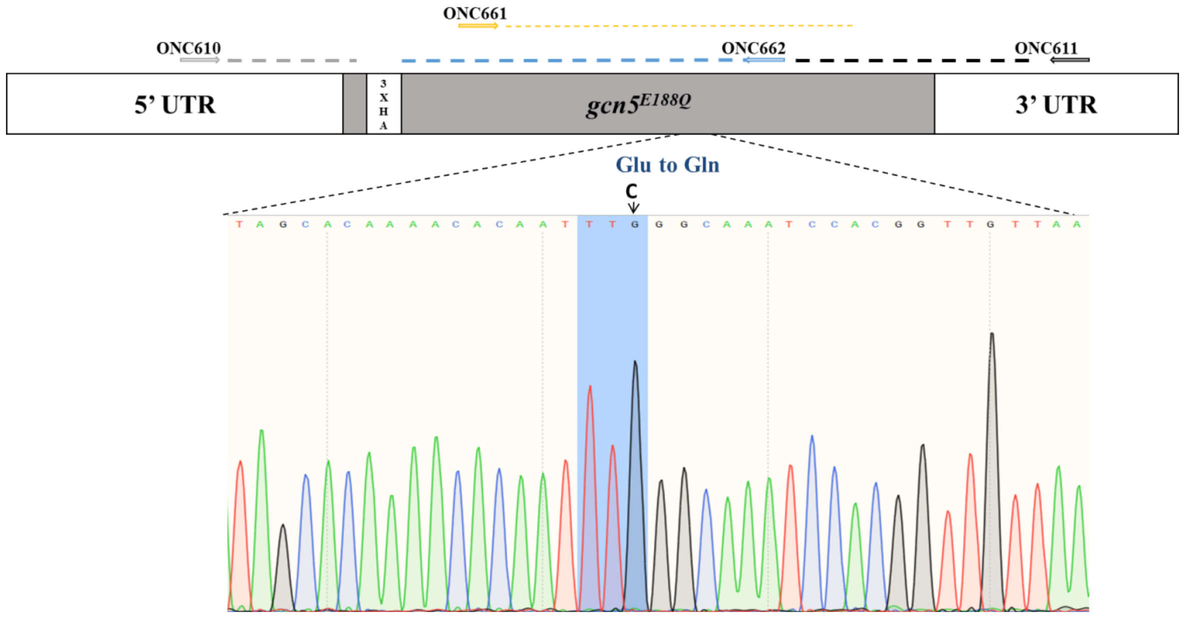


**b**

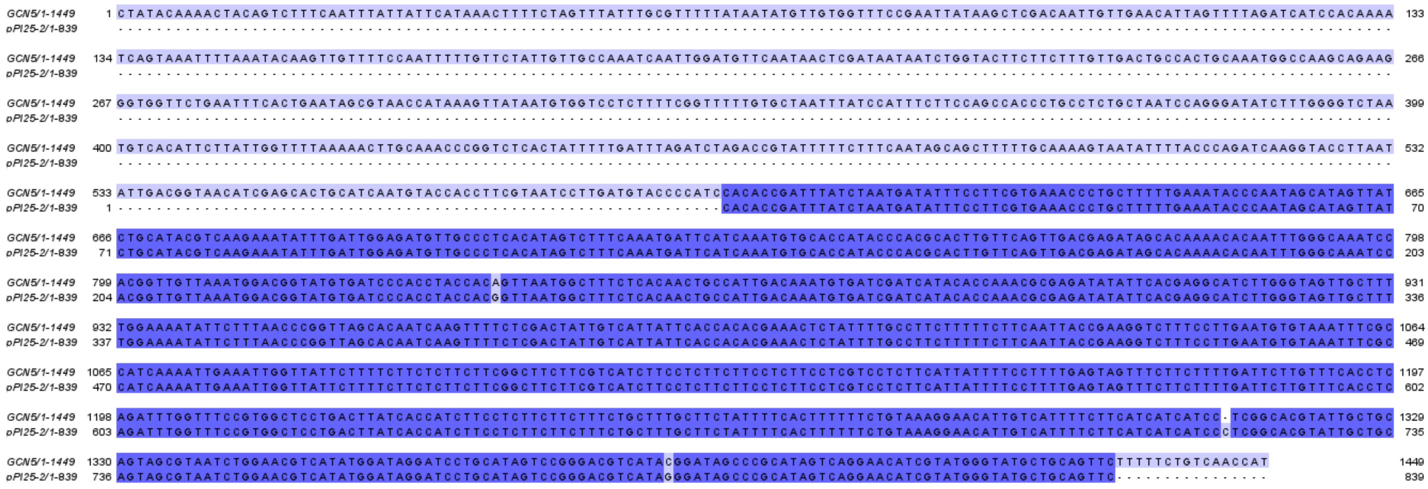


Nagar et al., Fig. S5

**a**

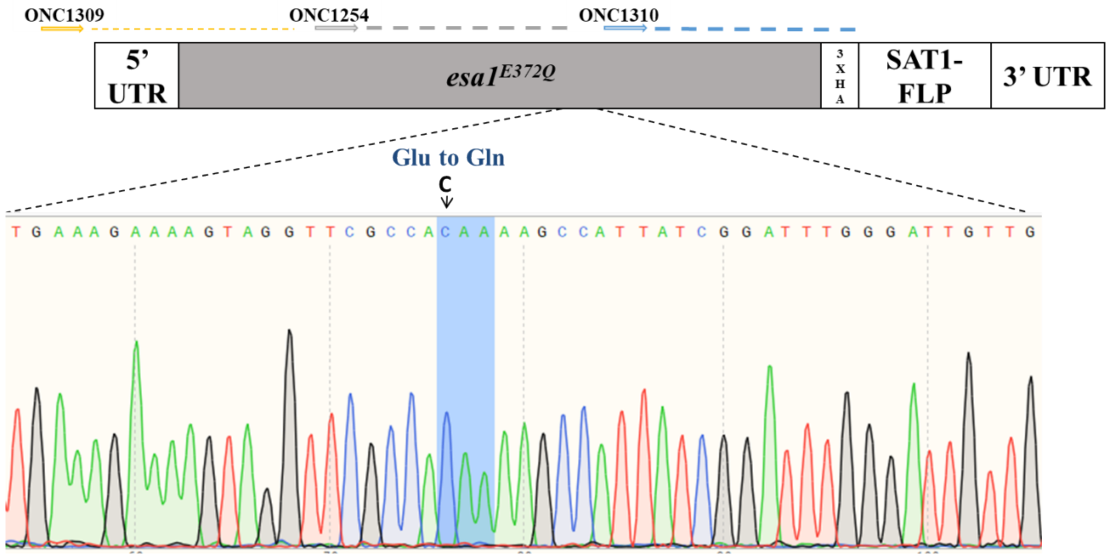


**b**

```
pPI22/1-756       1 TGCAGATGGATATAATGTGGCGTGTATTTTAACTCTACCCTGCTATCAAAAAAGAGGGTTTGGGAAATTGTTGATCCAGTTTTCATACATGCTAACGAAAGTTGAAAGAAAAGTAGGTTCGCCACAAAAGCCA 133
pPI22-51/1-802    1 ------------------------------------------------AAAAAGAGGGTTTGGGAA-TTGTTGATCCAGTTTTCATACATGCTAACGAAAGTTGAAAGAAAAGTAGGTTCGCCACAAAAGCCA 84

pPI22/1-756     134 TTATCGGATTTGGGATTGTTGTCATATCGTGCCTATTGGACAGACACATTGGTAAAGCTATTGGTAGAAAGAAACAGTCCTGCTTTATTCCGTAAGAATAATTCTCAGTTAGAGTACGATGAAGCTGAAAATG 266
pPI22-51/1-802   85 TTATCGGATTTGGGATTGTTGTCATATCGTGCCTATTGGACAGACACATTGGTAAAGCTATTGGTAGAAAGAAACAGTCCTGCTTTATTCCGTAAGAATAATTCTCAGTTAGAGTACGATGAAGCTGAAAATG 217

pPI22/1-756     267 GCAAAGACAGCTCTGCAACACCAACACCGGGGCCAGGTTCAAATGCCTCCCAATCATCTATATTGGCTAGTGCTGCTGCATCAAGGCTGGGACTTAATTCATCGCCTATTTTTTTCTAATGAAATTACCATTGA 399
pPI22-51/1-802  218 GCAAAGACAGCTCTGCAACACCAACACCGGGGCCAGGTTCAAATGCCTCCCAATCATCTATATTGGCTAGTGCTGCTGCATCAAGGCTGGGACTTAATTCATCGCCTATTTTTTTCTAATGAAATTACCATTGA 350

pPI22/1-756     400 GGACATTTCATCCATAACATGCATGACCACGACAGATATACTACACACATTAACCACATTACAGATGTTGCGATACTACAAGGGACAACACATAATTGTGTTAACCGATCAAATAATGGAATTGTATGAAAAA 532
pPI22-51/1-802  351 GGACATTTCATCCATAACATGCATGACCACGACAGATATACTACACACATTAACCACATTACAGATGTTGCGATACTACAAGGGACAACACATAATTGTGTTAACCGATCAAATAATGGAATTGTATGAAAAA 483

pPI22/1-756     533 CTAGTTAAAAAGGTAAAAGAGAAAAAGAAGCATGAGTTAAATCCAAAATTATTGCACTGGACACCACCTCTGTTCACTGCAAACCAATTACGGTTTGGGTGGTTAATTAACATCTTTTACCCATACGATGTTC 665
pPI22-51/1-802  484 CTAGTTAAAAAGGTAAAAGAGAAAAAGAAGCATGAGTTAAATCCAAAATTATTGCACTGGACACCACCTCTGTTCACTGCAAACCAATTACGGTTTGGGTGGTTAATTAACATCTTTTACCCATACGATGTTC 616

pPI22/1-756     666 CTGACTATGCGGGCTATCCGTATGACGTCCCGGACTATGCAGGATCCTATCCATATGACGTTCCAGATTACGCTGCTCAGTGCTGAGGCGC------------------------------------------ 756
pPI22-51/1-802  617 CTGACTATGCGGGCTATCCCTATGACGTCCCGGACTATGCAGGATCCTATCCATATGACGTTCCAGATTACGCTGCTCAGTGCTGAGGCGCGCCACTTCTAAATAAGCGAATTTCTTATGATTTATGATTTTT 749

pPI22/1-756         -----------------------------------------------------
pPI22-51/1-802  750 ATTATTAAATAAGTTATAAAAAAAAATAAGTGTATACAAATTTTAAAGTGACTC 802
```

Nagar et al., Fig. S6

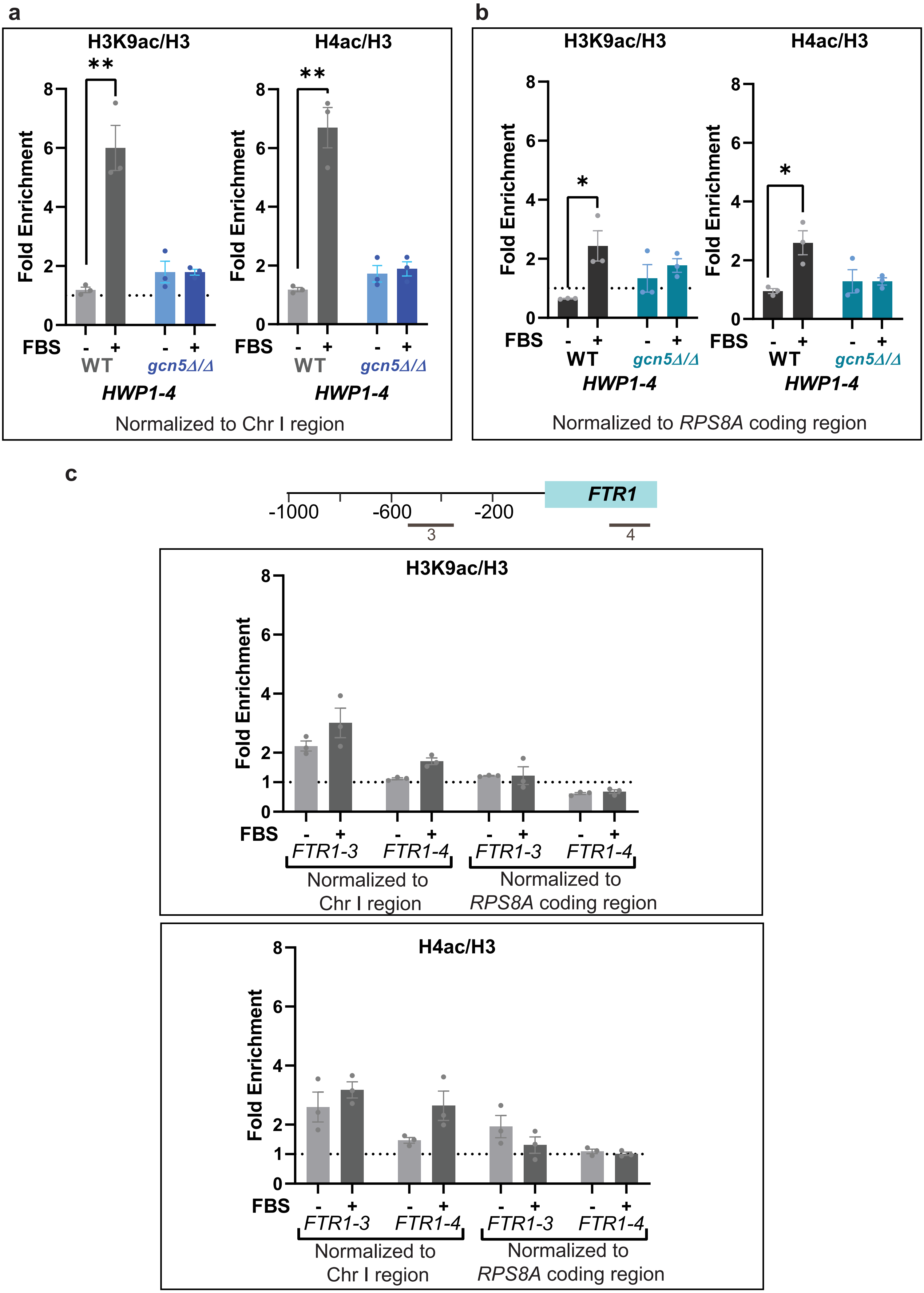


Nagar et al., Fig. S7
