## Supplementary material for "Histone acetylation by SAGA Complex but not by NuA4 Complex is Required for filamentation programme in *Candida albicans*": Legends for Supplental Figures

**Legends for Supplemental Figures**

**Fig. S1. Gcn5 and Esa1 are involved in regulating growth and filamentation of *C. albicans*.** Multiple sequence alignment of the (a) Gcn5 HAT domain and (b) Esa1 MYST domain amino acid sequences from *C. albicans, S. cerevisiae, Homo sapiens, Mus musculus* and *Drosophila melanogaster*. (c) Growth phenotype of Gcn5 catalytic dead mutants. The WT, *gcn5Δ/Δ*, and the catalytic dead mutant strains were grown in YPD, diluted and serial dilutions spotted on either YPD alone or on YPD with 10% FBS. The plates were incubated at 30°C or 37°C for 36h and image acquired. (d) Spot assay showing growth phenotype of WT and catalytically dead Esa1 under YPD and YPD with 10% FBS. The plates were incubated at 30°C or 37°C for 36h.

**Fig. S2. Histone H3K9ac and H4ac at hyphal gene promoters is mediated by Gcn5.**  Chromatin-level histone acetylation at (a) *HWP1* and (b) *ALS3* promoters using ChIP assay. Wild type (WT; SN95) and *gcn5Δ/Δ* strains were grown in YPD at 30°C or were induced for filamentation (YPD+10% FBS, 2h at 37°C), and equal amounts of chromatin extracts were used for immunoprecipitation with anti-H3K9ac, anti-H4ac (K5, K8, K12 and K16) antibodies or the control anti-H3 antibody. The immunoprecipitated DNA was analyzed by qPCR using primer pairs for different regions of *HWP1* and *ALS3* promoters or coding sequences, or a Chr I region as a nonspecific control. The data was quantified from three biological replicates each and the error bars represent SEM (n = 3). p-values ≤ 0.05 (∗), ≤0.01 (∗∗), and >0.05 (ns).

**Fig. S3. NuA4 complex recruitment is independent of Gcn5 at HSGs*.*** Epl1 recruitment at hyphal genes. Epl1-FLAG recruitment was analyzed by ChIP assay at (a) *HWP1* and (b) *ALS3* promoters in the wild-type strain (WT; SN95), PRI14 (*EPL1::FLAG_3_*) and PRI45 (*gcn5Δ/ΔEPL1::FLAG_3_*) strains. Cells were grown in YPD at 30°C or induced for filamentation (YPD+10% FBS, 2hr at 37°C) conditions, cross-linked with formaldehyde, harvested, lysed, and the chromatin extracts were prepared by sonication. Equal amount of chromatin extracts from strains were used for chromatin immunoprecipitation using anti-FLAG antibody. The enrichment of Epl1 was normalized to a nonspecific control Chr I region (ca21chr1_1573500–1574000) and WT (SN95) control. The quantification is based on three biological replicates and the error bars represent SEM (n = 3).  *p*-values ≤ 0.05 (∗), ≤0.01 (∗∗), and >0.05 (ns).

**Fig. S4. *Ca*Esa1 HAT activity is not required for H4 and H3K9 acetylation at HSGs.**

ChIP assay was used to measure the level of histone H3K9 acetylation and histone H4 acetylation at (a) *HWP1* and (b) *ALS3* promoters in wild type (WT; SN95) and *esa1^E372Q^*/*esa1^E372Q^* (PRI40) strains. Cells were grown in YPD at 30°C or induced for filamentation (YPD+10% FBS, 2h at 37°C), cross-linked with formaldehyde, harvested, lysed, and chromatin fragmented by sonication. Equal amounts of chromatin extracts were used for immunoprecipitation with anti-H3K9ac, anti-H4ac or, as control, anti-H3 antibodies and the immunoprecipitated DNA was analyzed by qPCR using primer pairs for *HWP1* and *ALS3* or a Chr I region as a nonspecific control. The data was quantified from three biological replicates each and the error bars represent SEM (n = 3). *p*-values ≤ 0.05 (∗), ≤0.01 (∗∗), and >0.05 (ns).

**Fig. S5. Construction of *HA_3_::gcn5^E188Q^* complemented strain expressed from native**

**promoter.** (a) Sequencing chromatograms showing incorporation of the E to Q mutation in

*GCN5*. (b) Alignment showing the sequencing result of glutamine substituted *GCN5* along

with the WT *GCN5* sequence.

**Fig. S6. Construction of *ESA1::HA_3_* and *esa1^E372Q^::HA_3_* plasmids.** (a) Schematic representation of plasmid pPI22 having *ESA1* and *esa1^E372Q^* in Ip30 vector. (b) Alignment showing the sequencing result of glutamine substituted *ESA1* along with the WT *ESA1* sequence.

**Fig. S7. Gcn5-mediated acetylation is linked to gene expression and filamentation.**  Histone H3K9 and H4 acetylation were assessed at *HWP1* (induced in FBS) and *FTR1* (uninduced in FBS) using ChIP assay. (a-b) The immunoprecipitated DNA was analyzed by qPCR using primer pairs for *HWP1-4* using either (a) Chr I region or (b) *RPS8A* coding region as normalization controls. (c) ChIP assay to assess H3K9 and H4 acetylation at *FTR1* gene. The immunoprecipitated DNA was analyzed by qPCR using primer pairs for different regions of *FTR1* promoter or ORF using Chr I region or *RPS8A* coding region as normalization controls. The data was quantified from three biological replicates each and the error bars represent SEM (n = 3). *p*-values ≤ 0.05 (∗), ≤0.01 (∗∗), and >0.05 (ns).
