## Supplemental Materials and Methods for "Histone acetylation by SAGA Complex but not by NuA4 Complex is Required for filamentation programme in *Candida albicans*"

**PLASMID AND STRAIN CONSTRUCTIONS**

**I. Construction of Catalytically dead Gcn5**

**Construction of HA_3_-tagged *GCN5.*** The HA**_3_** tag was amplified from Ip30 (Sinha et al., 2017) plasmid template using primers ONC1292 and ONC1293 having PstI sites on both primers. The 90 bp PCR product was digested with PstI. The plasmid pPI2 (Srivastav et al., 2021) contained multiple PstI sites, so the plasmid was truncated to remove the extra PstI sites in order to insert HA**_3_** tag inside *GCN5*. pPI12 plasmid was obtained by digesting pPI2-1 with HindIII, ligating and transforming into DH5α. pPI12 was digested with PstI and similarly digested PCR product was cloned into this vector. The clones were confirmed by PCR for orientation of HA**_3_** tag using primers ONC1292 and ONC662 which gave an amplified product of 893 bp. The positive clones which linearized upon NdeI digestion and gave a pop out of 451 bp upon BamHI digestion were named pPI13. The NAT^R^ cassette (excised from pPI2 by HindIII digestion) was reintegrated into HindIII digested pPI13 vector. The positive clones after confirmation with PCR using primers ONC611 and ONC661 which resulted in an amplification of 1558 bp product; and restriction digestion with BglII were named pPI15.

**Plasmid construction for FAE to AAA Site Directed Mutagenesis of *GCN5.*** Complementary primers incorporating the alanine substitutions (F186A, E188A) in place of conserved residues of interest were designed. The GST-*GCN5* vector was used as template for mutagenesis. The *GCN5* was amplified as two fragments with 53 bp overlapping region. ONC1256 and ONC464 amplified a fragment of 586 bp. Similarly, ONC1257 and ONC465 amplified a fragment of 814 bp. The two fragments were gel purified and used to amplify full length mutated *GCN5* by using equimolar ratio of both fragments. The PCR product and vector were digested with EcoRI and XhoI. The mutated *GCN5* was cloned into GST-*GCN5* vector and mutants were confirmed by sequencing with primer ON173. Clone pPI16-1 was found to contain the desired mutation.

The mutated *GCN5* was further sub-cloned into pPI13 by replacing the portion with mutated *GCN5* fragment of 865 bp excised from pPI16-1 by restriction digestion with BglII and PstI. The clones were confirmed by ORF specific PCR with ONC661 and ONC662 along with HA**_3_** tag orientation PCR using primers ONC1292 and ONC662. The construct was named pPI17. The NAT^R^ cassette was integrated into pPI17 by the same procedure as mentioned above to obtain pPI18. The pPI18 clones were confirmed by BglII digestion.

**Plasmid construction for E to Q Site Directed Mutagenesis of *GCN5.*** Complementary primers incorporating the alanine substitutions (E188Q) were designed. The pPI15-2 vector was used as template for mutagenesis. The *GCN5* sequence was amplified as two fragments with 23 bp overlapping region. ONC1312 and ONC610 were used to amplify a fragment of 998 bp. Similarly, ONC1311 and ONC611 amplified a fragment of 1130 bp. The two fragments were gel purified and used to amplify full length mutated *GCN5* by using equimolar ratio of both fragments. The PCR product and vector were digested with AatII and AflII. The mutated *GCN5* was cloned into similarly digested pPI15-2 vector and mutants were confirmed by HindIII digestion and sequencing (Fig.1). Clone pPI25-2-1 was found to contain the desired mutation by sequencing.

***C. albicans* integration of *HA_3_::GCN5*, *HA_3_::gcn5^FAE-AAA^* and *HA_3_::gcn5^E188Q^.*** The plasmids pPI15-2, pPI18-3 and pPI25-2-1 were linearized by digestion with PmeI and integrated in to KNC642 (*gcn5Δ/Δ)* strain and plated on 100µg/ml Nourseothricin containing

YPD plates. The colonies obtained were screened for integration by PCR using primers ONC661 and ONC662. The *HA_3_::GCN5* clones were named PRI20, *HA_3_::gcn5^FAE-AAA^* clones were named PRI21 and *HA_3_::gcn5^E188Q^* clones were named PRI33.

**II. Construction of Catalytically dead Esa1**

**Construction of HA_3_-tagged *ESA1.*** The Ip30 plasmid containing HA**_3_** tag with NAT^R^ cassette was used to clone *ESA1* along with 100bp of 5’UTR. The template of ~1700 bp was amplified from SN152 strain using primers ONC1295 and ONC1296 having PacI flanking sites. The Ip30 vector and PCR products were digested with PacI, ligated and transformed into DH5α. The clones were checked for correct orientation by PCR using ONC1293 and ONC1254 which gave a diagnostic band of 1310 bp. The positive clones were named pPI21.

**Site Directed Mutagenesis of *ESA1.*** Complementary primers incorporating the glutamine substitution in place of conserved glutamic acid residue were designed. The SN152 gDNA was used as template for mutagenesis. *ESA1* was amplified as two fragments with 50 bp overlapping region. ONC1297 and ONC1296 amplified a fragment of 542 bp. Similarly, ONC1298 and ONC1295 amplified a fragment of 1253 bp. The two fragments were gel purified and used to amplify full length mutated *ESA1*. The PCR product and Ip30 vector were digested with PacI. The mutated *ESA1* clones were named pPI22. The clones were further sequenced with primers ONC1254, ONC1309 and ONC1310 to confirm the complete sequence (Fig.2). Clone pPI22-51 was found to contain the desired mutation.

**Addition of 3’UTR to *ESA1::HA_3_* and *esa1^E372Q^*::*HA_3_* plasmid.** The 3’UTR region of *ESA1* was amplified with primers ONC1313 and ONC1314 containing ApaI and AflII sites respectively. The plasmids pPI21-28 and pPI22-51 were digested sequentially with AflII and ApaI, dephosphorylated and gel purified. The PCR amplified product of 1014 bp was digested similarly with AflII and ApaI and ligated to the vector. The positive clones were confirmed by PCR and digestion with HindIII. The pPI21 derivatives were named pPI26, and pPI22 derivatives were named pPI27.

***C. albicans* integration of *ESA1::HA_3_* and *esa1^E372Q^*::*HA_3_*.** The plasmids pPI26-1 and pPI27-15 were digested with NheI and AflII to obtain a linearized fragment of ~7Kb. The gel purified ~7Kb fragment was transformed into SN95 strain. The transformants were selected on 100µg/ml Nourseothricin containing YPD plates. The colonies obtained were screened for integration by PCR using primers ONC140 and ONC141 which gave a band of ~1000 bp. A second PCR with ONC1253 and ONC140 resulting in 2786 bp amplicon was used to detect the presence of cassette. The *ESA1*::*HA_3_* clones were named PRI35 and *esa1^E372Q^*::*HA_3_* clones were named PRI36.

The *SAT1* marker was recycled by growing PRI35 and PRI36 in presence of maltose and then screening for loss of marker. The clones thus obtained were named PRI37 (*ESA1::HA_3_*) and PRI38 (*esa1^E372Q^*::*HA_3_*). The strains were confirmed for loss of marker by nourseothricin cassette-specific PCR using primers ONC140 and ONC141. To construct homozygous *esa1^E372Q^*::*HA_3_* strain, the second copy of *esa1^E372Q^*::*HA_3_* was integrated by transforming the linearized plasmid cassette pPI27 resulting in clone PRI40.

To construct *gcn5Δ/Δ* and Esa1 catalytically dead strain, pPI27-15 was digested with NheI and AflII to obtain a linearized fragment of ~7Kb. The gel purified ~7Kb fragment was transformed into KNC642 *(gcn5Δ/Δ)* strain. The transformants were selected on 100µg/ml Nourseothricin containing YPD plates. The colonies obtained were screened for integration by PCR using primers ONC140 and ONC141. A second PCR with ONC1253 and ONC140 was used to detect the presence of cassette. The first allele integration clones were named PRI41. Further, to integrate both copies the SAT1 marker was recycled but we were unsuccessful in replacing both alleles of *ESA1*.
