## Supplemental Tables S1-S3 for "Histone acetylation by SAGA Complex but not by NuA4 Complex is Required for filamentation programme in *Candida albicans*"

**Table S1. List of Plasmids**

| **PLASMID** | **RELEVANT DESCRIPTION** | **SOURCE** |
| --- | --- | --- |
| pNIM1 | *C.a. SAT1 CaADH1* p*Tet-CaGFP* | (1) |
| pLITMUS28 | Cloning vector, Amp^R^ | NEB |
| pET28b (+) | T7 lac promoter with N- and C-terminal 6xHis tag | Novagen |
| pGEX-5X-3 | P_tac_promoter with N-terminal GST tag | GE Healthcare |
| pCIp10-*C.d. ARG4* | pCIp10 with *C.d. ARG4* in place of *C.a.URA3* | (2) |
| pHAH1 | HAH1 disruption cassette | (3) |
| Ip22 | TAP tag, *C.d. HIS1* | (2) |
| Ip27 | *SAT1FLP* cassette in pLITMUS28 | (2) |
| Ip30 | *HA_3_* tag, *SAT1FLP* cassette in pLITMUS28 | (2) |
| Ip50 | 6xHis-2xGly-3xFLAG tag, *ACT1*t,with *SAT1 FLP* | (2) |
| pPC46 | HA_3_ tag, *ADH1*t,*SAT1 FLP* in pLITMUS28 vector | This work |
| pPI1 | pCIp10 with *C.a. SAT1* in place of *C.a.URA3* | This work |
| pPI12 | pPI2 without NAT^R^ | This work |
| pPI13 | pPI12 with 3xHA tag in Ca*GCN5* | This work |
| pPI15 | pPI2 with 3xHA tag in Ca*GCN5* | This work |
| pPI16 | Ca*GCN5m* in BL21 | This work |
| pPI17 | pPI12 with 3xHA tag in Ca*GCN5^FAE^* | This work |
| pPI18 | pPI2 with 3xHA tag in Ca*GCN5^FAE^* | This work |
| pPI21 | Ip30 with *ESA1::HA_3_* | This work |
| pPI22 | Ip30 with *esa1^E372Q^::HA_3_* | This work |
| pPI23 | pPC68 with Ca*gcn5^FAE to AAA^ ::HA_3_* | This work |
| pPI24 | pPI2 with Ca*gcn5^E188Q^::HA_3_* | This work |
| pPI25 | pPI15 with Ca*gcn5^E188Q^::HA_3_* | This work |
| pPI26 | pPI21 with *ESA1* 3’UTR | This work |
| pPI27 | pPI22 with *ESA1* 3’UTR | This work |

| **STRAINS** | **RELEVANT GENOTYPE** | **SOURCE** |
| --- | --- | --- |
| SN95 | *arg4Δ/arg4Δ his1Δ/his1Δ URA3/ura3Δ::imm^434^ IRO1/iro1Δ::imm^434^* | (1) |
| KNC642 | *gcn5Δ::HAH1/gcn5Δ::HIS1* | (4) |
| TF156 | *efg1Δ::C.m.LEU2/ efg1Δ::C.d.HIS1* | (5) |
| ISC46 | SN95 *ADA2::TAP-C.d.HIS1/ADA2::TAP-C.m.LEU2* | (2) |
| PRI1 | *gcn5Δ::HAH1/gcn5Δ::HIS1<pPI2>* | (4) |
| PRI12 | ISC46 *EPL1::FLAG-SAT1 FLP* /*EPL1* | This work |
| PRI13 | ISC46 *EPL1::FLAG-FRT* */EPL1* | This work |
| PRI14 | ISC46 *EPL1::FLAG-FRT* / *EPL1::FLAG-SAT1 FLP* | This work |
| PRI15 | ISC46 *EPL1::FLAG-FRT* / *EPL1::FLAG-FRT* | This work |
| PRI20 | *gcn5Δ::HAH1/gcn5Δ::HIS1<pPI15>* | This work |
| PRI21 | *gcn5Δ::HAH1/gcn5Δ::HIS1<pPI18>* | This work |
| PRI23 | *gcn5Δ::HAH1/gcn5Δ::HIS1<pPI20>* | This work |
| PRI29 | *gcn5Δ::HAH1/gcn5Δ::HIS1<pPI23>* | This work |
| PRI30 | *gcn5Δ::HAH1/gcn5Δ::HIS1<pPI24>* | This work |
| PRI33 | *gcn5Δ::HAH1/gcn5Δ::HIS1<pPI25>* | This work |
| PRI35 | SN95 *ESA1::HA_3_-SAT1 FLP* /*ESA1* | This work |
| PRI36 | SN95 *esa1^E372Q^::HA_3_-SAT1 FLP* /*ESA1* | This work |
| PRI37 | SN95 *ESA1::HA-FRT* /*ESA1* | This work |
| PRI38 | SN95 *esa1^E372Q^::HA-FRT* /*ESA1* | This work |
| PRI39 | SN95 *esa1^E372Q^::HA-FRT* / *esa1^E372Q^::HA-FRT* | This work |
| PRI40 | SN95 *esa1^E372Q^::HA_3_-FRT* / *esa1^E372Q^::HA_3_-SAT1 FLP* | This work |
| PRI41 | *gcn5Δ::HAH1/gcn5Δ::HIS1 esa1^E372Q^::HA_3_-SAT1 FLP* /*ESA1* | This work |
| PRI42 | *gcn5Δ::HAH1/gcn5Δ::HIS1 EPL1::FLAG-SAT1 FLP* /*EPL1* | This work |
| PRI44 | *gcn5Δ::HAH1/gcn5Δ::HIS1 EPL1::FLAG-FRT* /*EPL1* | This work |
| PRI45 | *gcn5Δ::HAH1/gcn5Δ::HIS1 EPL1::FLAG-FRT* / *EPL1::FLAG-SAT1 FLP* | This work |

**Table S2. List of Strains**

| **OLIGO NUMBER** | **LENGTH** | **SEQUENCE (5’ to 3')** | **NOTES** |
| --- | --- | --- | --- |
| ON173 | 25 | GGG CTG GCA AGC CAG GTT TGG TGG T | pGEX-5X-3 sequencing primer |
| ONC114 | 18 | GGTGCCACTGATCCATTG | Position 61 to 78 within Ca*ARG4ORF* (Ca*ARG4*-F61) |
| ONC115 | 23 | GCCAACATATCCATAGTTAAAGC | Position 1108 to 1130(c) within *CaARG4ORF* (Ca*ARG4*-R1130) |
| ONC123 | 18 | GCCCTTCTGCCTGGAGTA | Diagnostic PCR primer within the non-repeat sequence of *HIS1* of pHAH |
| ONC140 | 24 | TCTTGGTGAGAACAGCGACCGAAA | Reverse primer for upstream split marker of SAT1 flipper |
| ONC141 | 24 | GGAGCGATAAGCGTGCTTCTGCCG | Fwd primer for downstream split marker of SAT1 flipper |
| ONC176 | 21 | TGCTCGTGTACCTGCTGTTTG | Fwd RT PCR primer for *CaSCR1* |
| ONC177 | 28 | ACACTGATTACTCTAGCGTTCAGAATTC | Rev RT PCR primer for *Ca SCR1* |
| ONC202 | 23 | GCCATTCACCAAGAATTTGACTT | Forward RT primer for *FTR1* ORF |
| ONC203 | 25 | ACAATTCATCCTTTTCAGCTTTGTT | Reverse RT primer for *FTR1* ORF |
| ONC305 | 28 | TACCAAGGAACACCAATATCTAGTCACT | Fwd RT ChIP PCR primer at Chr I Non-coding seq |
| ONC306 | 23 | GATCACTTTTGTCCTTGGCAGTC | Rev RT ChIP PCR primer at Chr I Non-coding seq |
| ONC431 | 23 | AGTCGAAAGAAAATTGGCTGCTA | Forward RT primer for *RPS8A* ORF |
| ONC432 | 24 | CAGAACCGAATTGAGAGTCAACAG | Reverse RT primer for *RPS8A* ORF |
| ONC464 | 30 | GAC GAA TTC CAT GGT TGA CAG AAA AAG AAC | 5’ primer for *GCN5* with EcoRI site |
| ONC465 | 31 | GCA CTC GAG TAC AAA ACT ACA GTC TTT CAA T | 3’ primer for GCN5 with XhoI site |
| ONC591 | 20 | GTCCATGGCCAACATGTGAA | Forward RT primer for *NRG1* |
| ONC592 | 23 | GTGTTGTTGTCTGTTGCGTTTGT | Reverse RT primer for *NRG1* |
| ONC 610 | 21 | TGTTGCCACTTGAGCACCACC | Up Check Forward primer for knock-out confirmation *GCN5* |
| ONC 611 | 23 | ATTGGCCACATCTTGTGAGTCGG | Down Check Reverse primer for knock-out confirmation *GCN5* |
| ONC 661 | 22 | GGTGATAAGTCAGGAGCCACGG | Forward *ORF* specific primer for *GCN5* |
| ONC662 | 22 | CGGTAACATCGAGCACTGCATC | Reverse *ORF* specific primer for *GCN5* |

**Table S3. List of Oligonucleotides**

| ONC911 | 25 | AATTATTATTCCCACCACACGACAC | Forward RT primer for *FTR1* promoter |
| --- | --- | --- | --- |
| ONC912 | 23 | GATTAGGCCCGATGTACATTGTC | Reverse RT primer for *FTR1* promoter |
| ONC1048 | 28 | ACCATCTGTAACACATCAAGGTCATATA | Forward RT primer for *BRG1* |
| ONC1049 | 20 | ACGATTTCCGGCTGCACTAT | Reverse RT primer for *BRG1* |
| ONC1052 | 20 | CCCACAACAACAACAGCAGG | Forward RT primer for *HWP1* (+188bp to +308bp) |
| ONC1053 | 22 | GGTTCTTGTGGTTGTTGTGGGT | Reverse RT primer for *HWP1* (+188bp to +308bp) |
| ONC1054 | 25 | TGGCTCCACTTACAAATCATAGTGA | Forward RT primer for *UME6* |
| ONC1055 | 24 | CAATCCTAGTCCCAACTCCAGATC | Reverse RT primer for *UME6* |
| ONC1088 | 30 | GTG GTC GAC ATG GTT GAC AGA AAA AGA ACT | Forward Primer for *GCN5* complementation |
| ONC1089 | 23 | GGA CCT CGA GTA CAA AAC TAC AG | Reverse primer for *GCN5* complementation |
| ONC1156 | 26 | CAC GAG CTC ATA ACT TGA AGA TCC CG | Forward primer for GCN5 wrt 1kb upstream of ATG |
| ONC1157 | 29 | GTC GAG CTC CAC TGA AGT AGA CTT CCC AC | Reverse primer for GCN5 wrt 500bp downstream of TAA |
| ONC1189 | 22 | CAAGAAATACAGGAAACCCTCC | Forward RT primer for *HWP1* |
| ONC1190 | 21 | CATATCGTATGTTTCAAAGAG | Reverse primer for *HWP1* |
| ONC1191 | 23 | ATCAGTCCACTAATTCCATCAAT | Forward RT primer for *HWP1* (-892bp to -739bp) |
| ONC1192 | 26 | AAGGACAAGAAAAATATGGATACAGT | Reverse primer for *HWP1* (-892bp to -739bp) |
| ONC1193 | 25 | TTTCTAACTCGACTAATGCACTTTA | Forward RT primer for *HWP1* (-617bp to -496bp) |
| ONC1194 | 27 | ATCCAAAGAATTATAGATGACTTGTTA | Reverse primer for *HWP1* (-617bp to -496bp) |
| ONC1195 | 23 | AGAGTTTTGGTAGGCTCATAATC | Forward RT primer for *HWP1* (-432bp to -311bp) |
| ONC1196 | 23 | CTAACTAACTTATCCCGGTTGTG | Reverse primer for *HWP1* (-432bp to -311bp) |
| ONC1197 | 22 | TCATAACTGGAATATTTTTGCT | Forward RT primer for *HWP1* |
| ONC1198 | 20 | TTAGTTAAACGCATCAGGAA | Reverse primer for *HWP1* |
| ONC1199 | 22 | CCACCACATGTATCGACAATGC | RT Forward primer for *EFG1* |
| ONC1200 | 20 | GCTCGAGGCGTTCAACGTAT | RT Reverse primer for *EFG1* |

| ONC1252 | 23 | GTACTATTCCCTTTCGGCTATCG | Forward Up Check primer for *ESA1* |
| --- | --- | --- | --- |
| ONC1253 | 22 | TCTGGACAATCTCCCTCTTTGG | Reverse Down Check primer for *ESA1* |
| ONC1254 | 20 | GTACCCCACAGCCAAGTGAT | Forward *ORF* specific primer for *ESA1* |
| ONC1255 | 20 | ATGGCTTTTCTGGCGAACCT | Reverse *ORF* specific primer for *ESA1* |
| ONC1256 | 53 | ACGAGATAGCACAAAACACAATAGCGGCAGCTCCACGGTTGTTAAATGGACGG | Forward primer for *GCN5* mutagenesis |
| ONC1257 | 53 | CCGTCCATTTAACAACCGTGGAGCTGCCGCTATTGTGTTTTGTGCTATCTCGT | Reverse primer for *GCN5* mutagenesis |
| ONC1258 | 81 | ATTAGTCCATCTACGCAATCGATTAGATTTGGGTCTATGTTGTTGAATAGAACACGTAAA CAT CAT CAC CAT CAT CAT GGT | Forward primer for *EPL1* FLAG tagging |
| ONC1260 | 20 | AACCAATAGGGCCGAAGAGT | Forward Up Check primer for *EPL1-FLAG* tag confirmation |
| ONC1261 | 20 | GACTGCAGCATCCAAATCAA | Reverse Down Check primer for *EPL1-FLAG* tag confirmation |
| ONC1276 | 80 | TATATACCCCCTTTGATTAGTAATCTCGCCACAGTTGCCTGCCGCGGAAAAATTCATTGAGAAGTTCCTATTCTCTAGAA | Reverse Primer for EPL1 FLAG tagging |
| ONC1280 | 22 | AACAATCCGAAGCAACGTAAAG | Forward RT primer for *ALS3* (-654bp to -476bp) |
| ONC1281 | 22 | CGGAAGATGTTGTTGCCAATAG | Reverse RT primer for *ALS3* (-654bp to -476bp) |
| ONC1282 | 17 | TTAGGTCGCTGGTTGCC | Forward RT primer for *ALS3* (-475bp to -318bp) |
| ONC1283 | 19 | GCGGACTCAGGGTTTGTAA | Reverse RT primer for *ALS3* (-475bp to -318bp) |
| ONC1284 | 21 | ATGCACGTTCATACTTCCAAA | Forward RT primer for *ALS3* (-217bp to -54bp) |
| ONC1285 | 23 | ACTATCAGACCTCAATTCAAGGG | Reverse RT primer for *ALS3* (-217bp to -54bp) |
| ONC1286 | 22 | TGACTTGGTCTAATGCTGCTAC | Forward RT primer for *ALS3* (+88bp to +271bp) |
| ONC1287 | 22 | CACCATGAGCAGTCAAATCAAC | Reverse RT primer for *ALS3* (+88bp to +271bp) |
| ONC1288 | 21 | GTATCACCTGTATCCGCCTTT | Forward RT primer for *HWP1* (-223bp to -29bp) |
| ONC1289 | 25 | GGAGATTCCTGTTGTGTTTGATATT | Reverse primer for *HWP1* (-223bp to -29bp) |
| ONC1290 | 21 | GGTCAAGGTGAAACAGAGGAA | Forward RT primer for *HWP1* (+78bp to +275bp) |
| ONC1291 | 21 | TCTTGTGGCTGTTGTGGATAG | Reverse primer for *HWP1* (+78bp to +275bp) |
| ONC1292 | 30 | TTT CTG CAG CAT ACC CAT ACG ATG TTC CTG | Forward primer for *GCN5* HA tag with PstI site |
| ONC1293 | 33 | AAACTGCAGTAGCGTAATCTGGAACGTCATATG | Reverse primer for *GCN5* HA tag with PstI site |
| ONC1295 | 37 | TTTTTAATTAATGTACAGAGACAAAATCAATACATCA | Forward primer for *ESA1* HA tag with PacI and BsrGI site to clone in pLIT28-HA-SAT1 (pPC46) |
| ONC1296 | 30 | TTTTTAATTAACCACCCAAACCGTAATTGG | Reverse primer for *ESA1* HA tag with PacI site to clone in pLIT28-HA-SAT1(pPC46) |
| ONC1297 | 50 | GTT GAA AGA AAA GTA GGT TCG CCA CAA AAG CCA TTA TCG GAT TTG GGA TT | Forward primer for *ESA1* mutagenesis |
| ONC1298 | 50 | AATCCCAAATCCGATAATGGCTTTTGTGGCGAACCTACTTTTCTTTCAAC | Reverse primer for *ESA1* mutagenesis |
| ONC1309 | 19 | TAG GTG AGC TCT GGT ACC C | pLIT28-HA-SAT1 |
| ONC1310 | 22 | TGC AGA TGG ATA TAA TGT GGC G | *ESA1* ORF specific primer |
| ONC1311 | 23 | CCG TGG ATT TGC CCA AAT TGT GT | Forward primer for *GCN5* E to Q mutagenesis |
| ONC1312 | 23 | ACA CAA TTT GGG CAA ATC CAC GG | Reverse primer for *GCN5* E to Q mutagenesis |
| ONC1313 | 30 | GCA GGG CCC TAT ATT TAA TAG GAT GTT CCC | Forward primer for *ESA1* 3'UTR with ApaI site |
| ONC1314 | 29 | AAA CTT AAG GGC TGG CCG ACT CTA CAA TA | Reverse primer for *ESA1* 3'UTR with AflII site |
| ONC1315 | 20 | AAA TTG CGG ATA AAC TTG CC | Forward primer for *HWP1* (-1249bp to -1098bp) |
| ONC1316 | 20 | ATG GTT AGG CAA CTC TTA CC | Reverse primer for *HWP1* (-1249bp to -1098bp) |
| ONC1317 | 23 | GTG ATG CTG GAT CTA ACG GTA TT | Forward RT primer for *ALS3* (+952bp to +1049bp) |
| ONC1318 | 22 | CGG TTA GGA TCG AAT GGT AAG G | Reverse RT primer for *ALS3* (+952bp to +1049bp) |
| ONC1319 | 20 | TTC CAG CTA CCA CTC CAA AC | Forward RT primer for *HWP1* (+703bp to +801bp) |
| ONC1320 | 23 | TCC AGA AGT AAC TGG AAC AGA AC | Reverse primer for *HWP1* (+703bp to +801bp) |
